# Lipid nanoparticle protein coronas arise through lipoprotein fusion rather than shell-like adsorption

**DOI:** 10.64898/2025.12.21.695162

**Authors:** Shaun Grumelot, Naseeha Mohammed, Ghafar Yerima, Jorge Colonrosado, Seyed Amirhossein Sadeghi, Fei Fang, Kylie Hilsen, Brooke Shango, Amir Ata Saei, Amanda M. Murray, Michael J. Mitchell, Babak Borhan, Liangliang Sun, Hojatollah Vali, Mohammad R. K. Mofrad, Kathryn A. Whitehead, Morteza Mahmoudi

## Abstract

The protein corona influences the *in vivo* biodistribution of ionizable lipid nanoparticles (LNPs) in nucleic acid delivery, yet its structural architecture remains poorly defined. Using cryo-transmission electron microscopy, we visualized LNP-protein interactions in their native state. We show that, unlike the discrete “fuzzy” shells observed on hard nanoparticles, LNPs displayed no peripheral protein shell. Instead, controlled incubation and competitive “dual-particle” assays, supported by molecular dynamics simulations, indicate that LNP membranes undergo localized thickening and electron-dense remodeling consistent with lipoprotein integration rather than surface adsorption. Similar features were observed in extracellular vesicles, suggesting this behavior is shared among lipid-based carriers, and proteomic analysis identified apolipoproteins as the dominant associated proteins. Together, these findings support a model in which the biological identity of LNPs arises through membrane remodeling rather than shell-like adsorption, and provide a framework for the rational design of targeted nanomedicines.

**TOC Graphic:** 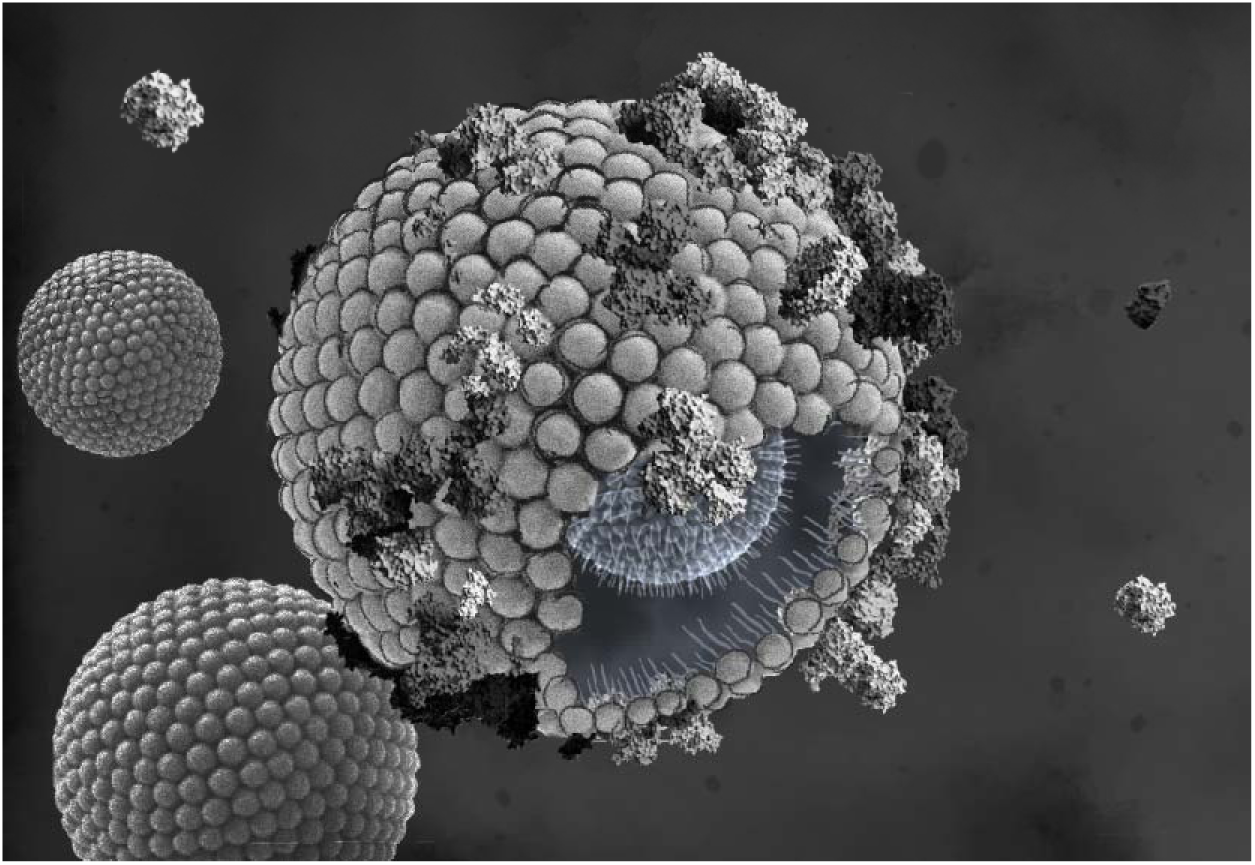

## Introduction

Upon physiological exposure via systemic administration, nanoparticles rapidly acquire a dynamic, protein-rich interfacial layer often described as the “protein corona”.^1, 2^ This complex layer, predominantly composed of plasma proteins, is a key determinant of nanoparticle biodistribution, cellular uptake, immune recognition, and ultimately, therapeutic efficacy.^3^ For hard nanoparticles such as those made from gold or polystyrene, this corona manifests as a stable, shell-like, “fuzzy” protein layer adsorbed to the particle surface, as visualized by cryo-transmission electron microscopy (cryo-TEM).^4, 5^ This concept has guided many strategies aimed at predicting and controlling the biological fate of nanoparticles.

In recent years, these strategies have been increasingly used to study ionizable lipid nanoparticles (LNPs), given their transformative role in the recent clinical breakthroughs of mRNA vaccines and gene therapies.^6, 7^ Biochemical characterization using mass spectrometry has identified proteins enriched on LNP surfaces after plasma exposure, including apolipoproteins and other plasma proteins, and demonstrated that these protein associations affect LNP cellular uptake, intracellular trafficking, and efficacy.^8^ Despite this functional evidence and compositional characterization, the molecular architecture of protein-LNP interaction – that is, how proteins organize on LNP surfaces – remains poorly understood. Many studies implicitly assume that LNPs, like hard nanoparticles, acquire conventional shell-like protein coronas. Nonetheless, direct visualization in the native, hydrated state has been limited.

Fortunately, recent advances in cryo-TEM have enabled this. For hard nanoparticles, cryo-TEM analyses reveal a distinct protein shell surrounding the particle.^4, 5^ However, analogous high-resolution studies of protein coronas on soft nanoparticles (e.g., LNPs) are lacking, leaving their coronal architecture unresolved. This study addresses this gap by combining cryo-TEM visualization of LNP protein associations and mass spectrometry to correlate structural observations with protein composition.

Although biochemical studies have identified proteins associated with LNP surfaces, only a few investigations have directly visualized LNPs that have been exposed to human serum/plasma using cryo-TEM. These investigations, which focused primarily on LNP structure, contain images that do not show a shell-like corona like those observed on hard nanoparticles.^8^ To investigate whether we could visualize protein coronas on LNPs and corroborate these prior observations, we employed cryo-TEM imaging with improved sample preparation methods. A critical challenge in LNP corona studies is contamination from extracellular vesicles (EVs), which share structural and compositional similarities with LNPs and can confound interpretation. ^9^ We therefore developed a workflow combining EV depletion, size exclusion chromatography purification, and cryo-TEM visualization. This approach reduces EV contamination and enables direct assessment of protein organization on LNP surfaces.

## Results and discussion

The study design is summarized in **Fig. 1A**. To minimize co-isolation of EVs with LNPs, we depleted marker-positive EVs, including exosomes and microvesicles, from human plasma immunomagnetic capture, binding EVs to magnetic MACSPlex multiplex beads and removing them with an external magnetic separator. The captured EVs were then characterized by flow cytometry, which reads 37 EV surface epitopes (**Fig. 1B**). For example, well-established exosome markers CD9, CD63, and CD81^10^ were successfully identified, confirming effective isolation of exosomes from human plasma. We subsequently performed liquid chromatography-tandem mass spectrometry (LC-MS/MS) on the isolated EVs and identified 166 proteins, establishing an EV reference proteome. We then used this reference to screen the LNP fractions, where these EV-marker proteins were detected at negligible levels, indicating that the LNP samples were not contaminated by co-isolated EVs.

**Fig. 1.**
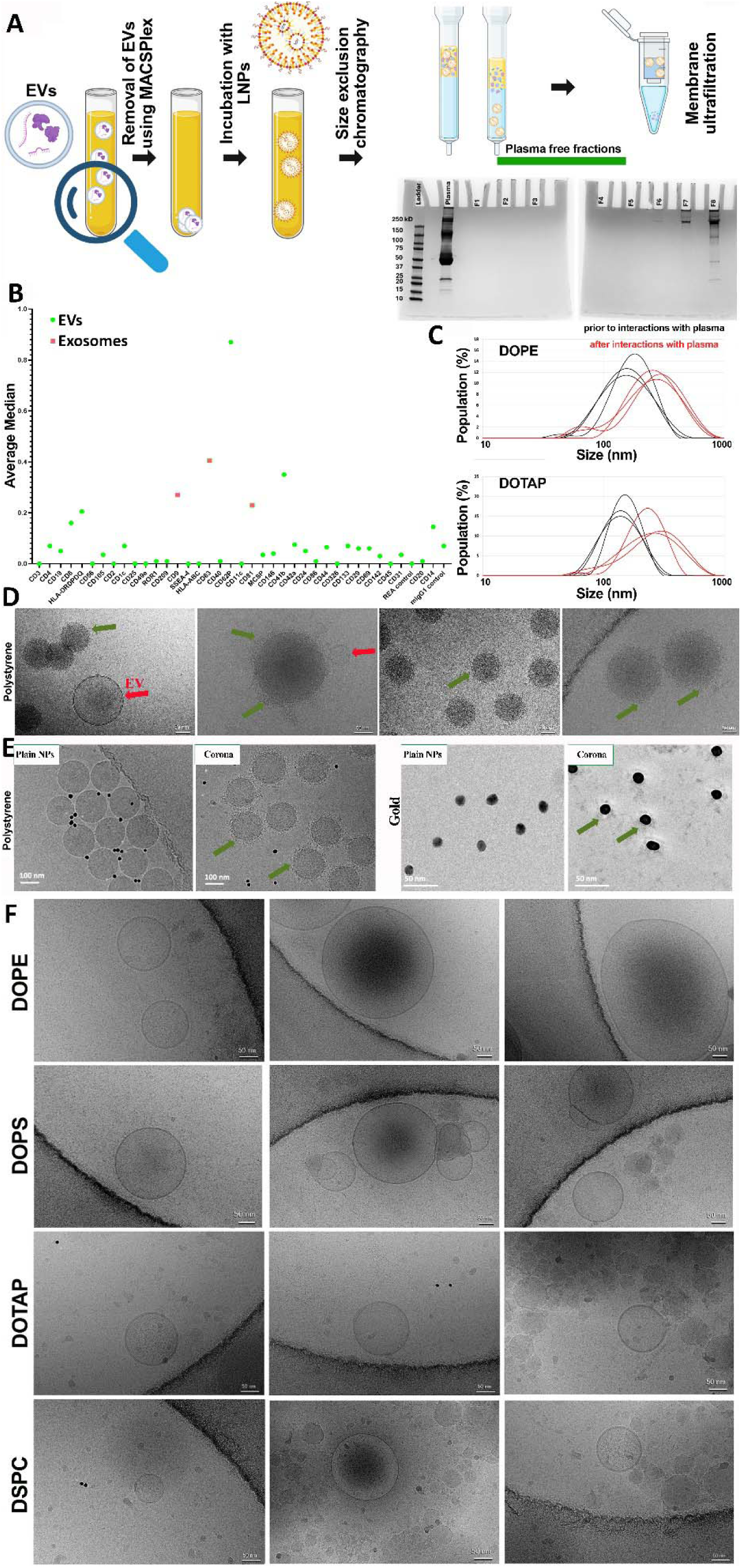
LNPs lack a conventional protein corona shell. **A)** Schematic of the study design: human plasma was first depleted of EVs; the EV-depleted plasma was then incubated with LNPs. The mixtures were passed through size exclusion chromatography (Sepharose CL-4B column), and fractions 2–6—containing LNPs (larger LNP particles pass through the column faster than smaller plasma proteins) with minimal free or excess plasma proteins, as confirmed by gel electrophoresis of collected plasma alone in various fractions—were collected. These LNPs underwent ultrafiltration using Vivaspin Columns (MWCO 1,000,000) to remove unbound proteins before mass spectrometry analysis. **B)** MACSPlex analysis showed expression of 37 surface epitopes on plasma-derived EVs, including exosome markers (CD9, CD63, CD81). **C)** Representative dynamic light scattering (DLS) results demonstrated an increase in LNP size after plasma exposure and fraction collection. **D)** Cryo-TEM images of polystyrene nanoparticles (100 nm and 200 nm) display characteristic fuzzy protein coronas (green arrows), while EVs lack such structures (red arrows). These findings indicate minimal or absent protein corona formation on EVs, significantly contrasting with the well-documented corona formation on polystyrene nanoparticles. **E)** Cryo-TEM images of polystyrene (100 nm) and gold (15 nm) nanoparticles before and after formation of protein corona. **F)** Cryo-TEM images reveal that LNPs do not form peripheral fuzzy protein shells like those observed on hard nanoparticles. Instead, some LNPs show electron-dense structures, consistent with lipoprotein particles, in close association with or fused into LNP membranes (see Fig. 2 for more details). This morphology was consistent across the four LNP formulations. Panel **E** is reproduced with permission from reference^5^, Copyright [2021] [Nature Publishing Group].

Next, we generated LNPs formulated with the ionizable lipidoid 306O_i10_, cholesterol, one of four helper lipids (DOPE, DSPC, DOPS, or DOTAP), and PEG-lipid.^11^ LNP formulation details and characterization data (**Table S1**) are available in Supplementary Information (**SI**). Each LNP was incubated with EV-depleted human plasma, and corona-bearing LNPs were then isolated using a standardized column separation method (see **Fig. 1A**).^12^ MACSPlex analysis of the collected plasma-derived EVs confirmed the successful collection of EVs from plasma (**Fig. 1B**). Dynamic light scattering (DLS) measurements before and after plasma incubation revealed increases in LNP hydrodynamic size upon interaction with plasma proteins (**Fig. 1C**). Subsequently, we employed cryo-TEM to image proteins in their native state on the surface of nanoparticles. Together with prior EV depletion and column purification, this approach permits direct observation of the LNP-protein interface while minimizing impurities from EVs and protein aggregates.^4, 5^

As a positive control for the formation of a typical protein corona on hard nanoparticles, we incubated polystyrene nanoparticles (100 nm and 200 nm) with human plasma, isolated the corona coated particles and imaged them. As expected, and consistent with prior reports,^4, 5^ these controls displayed a protein shell surrounding the particle surface. Interestingly, in a couple of images, we also identified EVs (**Fig. 1D**) that did not exhibit a discernable protein corona (**Fig. 1D**).

To further enhance the comparison of the fuzzy protein corona formation on nanoparticles, **Fig. 1E** presents polystyrene and gold nanoparticles before and after protein corona formation. Both nanoparticle types exhibit the development of a classic protein corona shell. Interestingly, even gold nanoparticles, which possess a high atomic mass and exhibit significant electron beam absorption, still reveal the presence of the protein corona shell. However, the intricate details of the protein features are less discernible in gold nanoparticles compared to the polystyrene nanoparticles, which have lower atomic mass and electron beam absorption. Surprisingly, and similar to EVs, cryo-TEM analysis of all four LNP formulations after plasma incubation revealed a structural organization distinct from classical protein coronas observed on hard nanoparticles (**Fig. 1F**). These observations indicate that lipid-based particles, such as LNPs and EVs may engage proteins through different structural mechanisms than hard nanoparticles.

Although most studies of LNP protein coronas have not employed cryo-TEM for direct visualization, one report^8^ does provide images consistent with our findings. In contrast to LNPs, cryo-TEM has been extensively used to study EV-EV and EV-lipoprotein interactions.^13,14^ In agreement with our observations for LNPs, the classic protein corona morphology (shell-like, fuzzy architecture) was not detected on the surface of various types of EVs isolated from different biological fluids, including human plasma.^13–15^ We also reviewed the literature on cryo-TEM analyses of EVs from human plasma and other biological fluids, which corroborates this lack of detectable protein corona formation on their surfaces.^16^ Taken together with our data, these findings suggest that lipid-based nanoparticles may not assemble a classical, well-defined protein corona structure (**Fig. 2A** and **B**), thereby challenging prevailing assumptions about lipid-protein interactions in biological media. Images of LNPs also revealed formation of electron-dense semi-spherical structures that fused into the LNP membranes. This morphology was seen across the four LNP formulations. These structures were frequently observed in EVs and various studies revealed that they are composed of lipoproteins.^13^

**Fig. 2.**
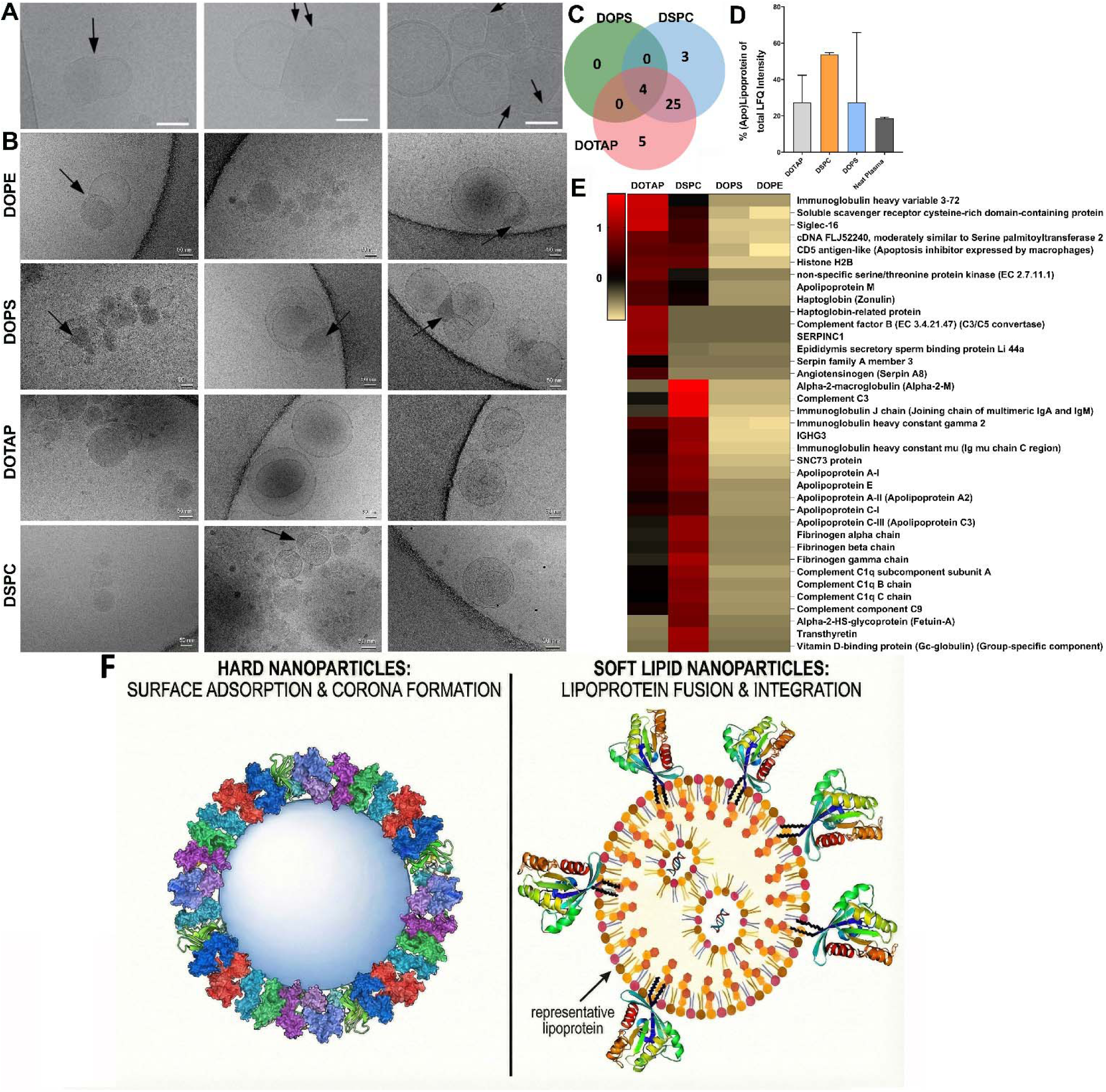
Cryo-TEM images showing the association and possible fusion of lipoproteins and other proteins with the surfaces of LNPs. **A)** Representative cryo-TEM images of EVs isolated from biological fluids, showing the absence of a conventional protein corona. Notably, these images also reveal binding and fusion events between EVs and lipoproteins, indicated by black arrowheads (scale bar = 50 nm; figures adapted with permission from Ref.^13^). **B)** Representative cryo-TEM images of various types of LNPs showing electron-dense structures associated with, and in some cases apparently integrated into, the LNP membrane (black arrowheads), consistent with lipoproteins rather than a peripheral protein shell. Alternatively, the black arrowheads may represent blebs, or a combination of blebs and fused lipoproteins. These findings suggest a role for either fusion-mediated interactions, protein interactions with blebs, or a combination thereof, rather than conventional corona formation. **C)** Venn diagram representing overlapping protein groups identified across all samples. Protein groups included in this analysis contained valid label-free quantification (LFQ) intensity values. Overlaps indicate proteins commonly detected between NPs, reflecting shared and unique components of the protein corona composition. **D)** Percentage contribution of (Apo)lipoproteins to total LFQ intensity in each sample. (Apo)lipoprotein levels in neat plasma were included as a control for baseline comparison. The bar graph shows the relative proportion of (Apo)lipoproteins compared with the total detected proteins across LNPs (n=6 for each LNP type). **E)** Heatmap of 37 statistically significant protein groups across samples. Rows represent protein groups filtered to include a minimum of three valid LFQ intensity values in at least one group per sample. ANOVA analysis identified proteins showing significant abundance differences. LFQ intensities were Z-score normalized, where red indicates higher and beige indicates lower relative abundance. **F)** Schematic showing the distinct mechanisms of protein-nanoparticle interaction. Hard nanoparticles (left panel) recruit proteins into organized peripheral shells through surface adsorption. Soft lipid-based nanoparticles (right panel) engage proteins primarily through lipoprotein/lipidated-protein fusion and integration into the bilayer.

To determine the composition of the structures associated with LNPs in our cryo-TEM images, we employed LC-MS/MS bottom-up proteomics on the collected LNPs after contact with plasma proteins. The results revealed a substantial (apo)lipoprotein signature within the LNP-associated fractions (**Fig. 2C-E**). Because lipoproteins are similar in size and density to LNPs, they co-elute during size exclusion chromatography, so their abundance in these fractions identifies the associated species but does not by itself separate integrated from co-isolated particles. Apolipoproteins accounted for a mean of ∼25–55% of total LFQ intensity across the LNP fractions (varying by LNP formulation), compared with approximately 18% in neat plasma processed in parallel (**Fig. 2D**), indicating selective association. The structural basis for integration, rather than adsorption or co-elution, is established by the cryo-TEM morphology and the simulations described below.

Across the four LNP formulations, each prepared in two independent syntheses and analyzed in three separate LC-MS/MS runs per batch, we identified 124 unique protein groups, counting each protein once across formulations. Individual formulations contributed 1 (DOPE), 7 (DOPS), 61 (DOTAP), and 71 (DSPC), with many proteins shared between DOTAP and DSPC. Of the 124, 37 protein groups were reproducibly detected across replicates (DOPE, 0; DOPS, 4; DOTAP, 34; DSPC, 32; **Fig. 2C**). DOPE LNPs yielded few or no reproducibly detected proteins, so the apolipoprotein signature rests on the DOTAP and DSPC formulations rather than uniformly across all four, and we interpret the DOPE and DOPS groups with caution given their low absolute counts. The large standard deviation for the DOPS and DOTAP in **Fig. 2D** reflects sparse, near-detection-limit signal rather than wide biological spread. The raw proteomics data revealed abundant charge-1 lipid-related ions that reduce peptide ionization efficiency through ion suppression. Because the degree of this interference differed between runs, the measured apolipoprotein LFQ intensities also varied across replicates. In groups like DOPS, where only a few proteins were detected, this makes the standard deviation especially sensitive to small fluctuations.

To assess whether PEG content alters protein association, we compared DOPE LNPs prepared with 0%, 1.5%, and 2.5% PEG. The 0% and 1.5% formulations showed notable aggregation, while the number and identities of associated proteins did not differ across the three formulations. Because DOPE LNPs returned few reproducibly detected proteins overall, we do not draw a strong conclusion regarding PEG dependence.

Using lipoprotein association fluorometry (LAF), a fluorescent assay that utilizes lipophilic indocarbocyanine dyes, a recent study demonstrated that lipoproteins associate with various types of EVs.^13^ Utilizing LAF with cryo-TEM, Wolfram and colleagues^13^ showed that EVs derived from human, nematode, and bacterial sources bind to multiple lipoprotein classes, including very-low-density lipoprotein (VLDL) and low-density lipoprotein (LDL). Their work, consistent with many other studies (e.g.,^17^), further indicates that lipoproteins can not only bind to, but also fuse with, different EV types. In line with these findings, we observed substantial lipoprotein-LNP attachment and structures consistent with membrane fusion in human plasma (**Fig. 2B**), mirroring phenomena frequently reported for EVs.^13, 16^ Importantly, our corona-LNPs showed no detectable EV surface-marker signal by MACSPlex analysis. Beyond lipoprotein-LNP attachment and membrane fusion, the multicompartment LNPs observed in **Fig. 2B** could also arise from “blebs” (i.e., distinctive mRNA-rich structures),^18, 19^ which are reported in literature for a subset of LNPs. These blebs, which separate from the LNP lipid bilayer, may be particularly prone to protein interactions.

Given that the association of apolipoprotein E (ApoE) is widely recognized as a primary driver of LNP biodistribution and cellular uptake within the liver, we performed controlled incubation studies with purified ApoE4 (one of the three common isoforms of ApoE) and DOTAP LNPs to mechanistically characterize this lipid-protein interface (**Fig. 3**). Cryo-TEM imaging of the ApoE4 control revealed a dense, granular distribution of individual protein molecules. The DOTAP control LNPs exhibited a characteristic clean, well-defined unilamellar lipid bilayer with no surface adsorbents.

**Fig. 3.**
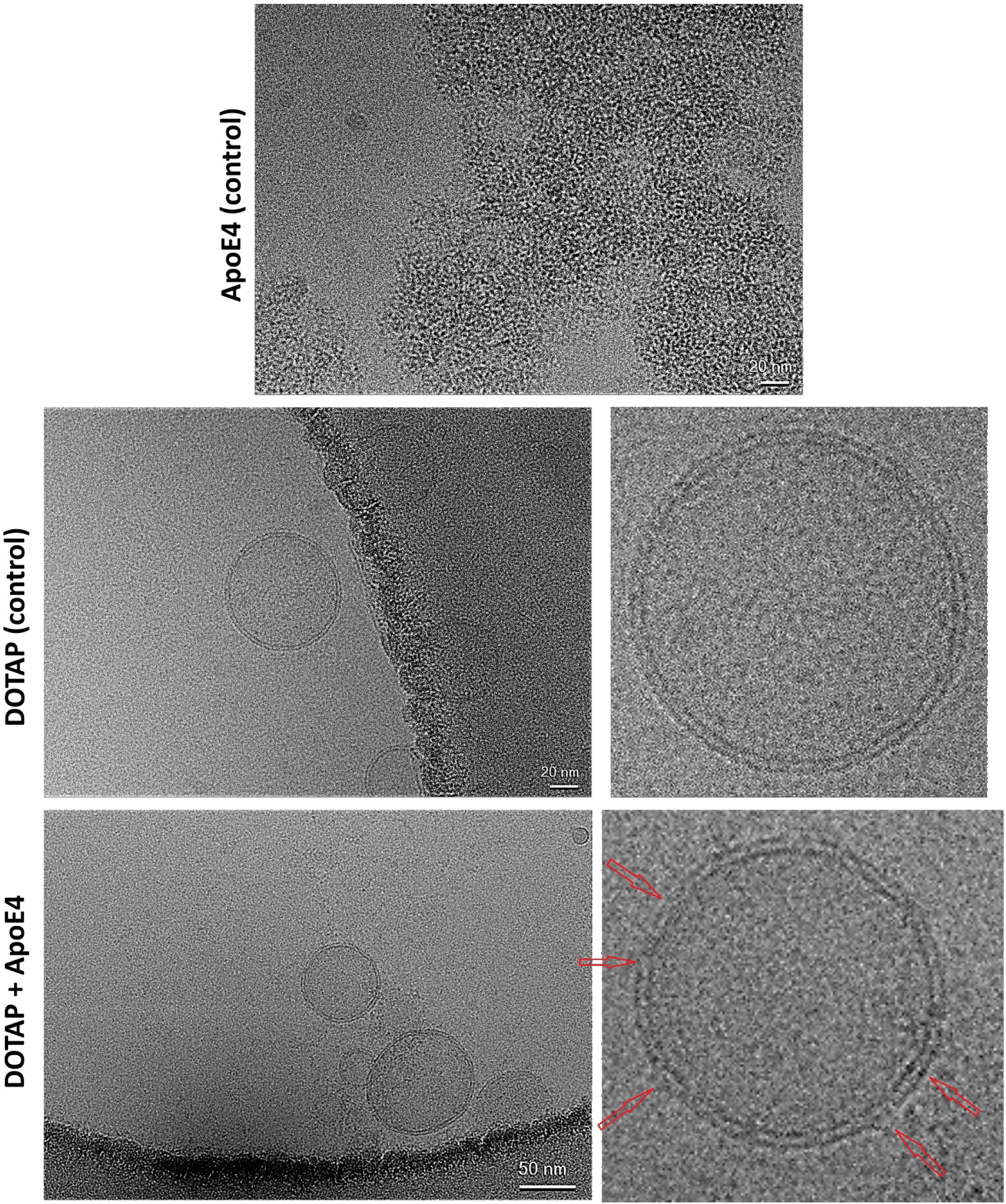
Cryo-TEM analysis of ApoE4–LNP interactions reveals membrane remodeling consistent with fusion-like behavior. Cryo-TEM images of (top) ApoE4 alone, (middle) DOTAP LNPs, and (bottom) DOTAP LNPs following incubation with ApoE4. ApoE4 alone forms amorphous, non-vesicular assemblies without defined bilayer structures. Control DOTAP LNPs display spherical morphology with smooth, well-defined lipid bilayers. In contrast, ApoE4-incubated LNPs retain overall vesicular integrity but exhibit localized membrane irregularities, including bilayer thickening and asymmetric density distributions (red arrows). Notably, no continuous outer protein layer is observed, arguing against formation of a classical protein corona. Instead, these features are consistent with membrane insertion and fusion-like interactions, suggesting partial integration of ApoE4 into the LNP lipid bilayer.

Cryo-TEM analysis of LNPs incubated with ApoE4 reveals that ApoE4 does not form a conventional, uniform protein corona around LNPs (**Fig. 3**). Instead, ApoE4 induces localized membrane remodeling, manifested as bilayer thickening and asymmetric electron density distributions while preserving overall vesicular integrity. The red arrows in **Fig. 3** highlight specific regions where the LNP membrane exhibits localized thickening and increased electron density, consistent with the hydrophobic insertion and diffusion of ApoE4 into the lipid leaflet. These features suggest that the LNP surface is dynamically remodeled by apolipoproteins. These features are inconsistent with simple surface adsorption and instead suggest membrane insertion and fusion-like interactions, likely mediated by the amphipathic α-helical domains of ApoE4 that enable strong lipid binding.

This behavior aligns with ApoE4’s physiological role in lipoprotein remodeling, where it dynamically associates with lipid surfaces and can partially embed within lipid bilayers. Such integration may facilitate lipid–protein mixing and alter membrane packing, potentially influencing LNP stability, surface properties, and interactions with biological systems. Importantly, this mechanism provides a conceptual shift from the traditional protein corona paradigm toward a model of protein–lipid co-assembly, with significant implications for understanding nanoparticle identity *in vivo*. While cryo-TEM alone cannot definitively resolve the extent of molecular insertion or membrane fusion, the observed structural perturbations strongly support a model of partial ApoE4 incorporation into the LNP membrane, warranting further investigation using complementary cryo-electron tomography and/or computational techniques.^5, 20, 21^

Our findings align with prior studies indicating that ApoE disrupts the integrity of lipid bilayers.^22–24^ Studies revealed a coordinated, multi-step mechanism.^22^ The protein first targets membrane defects, then the N-terminal bundle unfolds via interactions with the highly hydrophobic C-terminal domain, producing a large conformational change that exposes the protein’s buried hydrophobic core to the lipid surface. Next, ApoE intercalates into the bilayer and stabilizes edges: its hydrophobic faces insert into the membrane, isolate lipid patches, and wrap around the perimeter like a belt to shield exposed regions from water, effectively forming nanolipoprotein-like structures. Finally, ApoE drives component redistribution, altering internal lipid packing and extracting lipids from the LNP until the original bilayer loses structural integrity.^22, 25, 26^

To test whether the membrane remodeling we observed is specific to LNP architecture and not a general property of ApoE4, we performed a competitive incubation study. First, we confirmed that ApoE4 forms a conventional “fuzzy” protein shell on hard, non-lipid surfaces by incubating it with polystyrene NPs. Cryo-TEM analysis of polystyrene NPs alone incubated with ApoE4 clearly revealed a well-defined, robust peripheral protein corona shell (**Fig. 4A**), consistent with classical adsorption models on solid-core NPs.

**Fig. 4.**
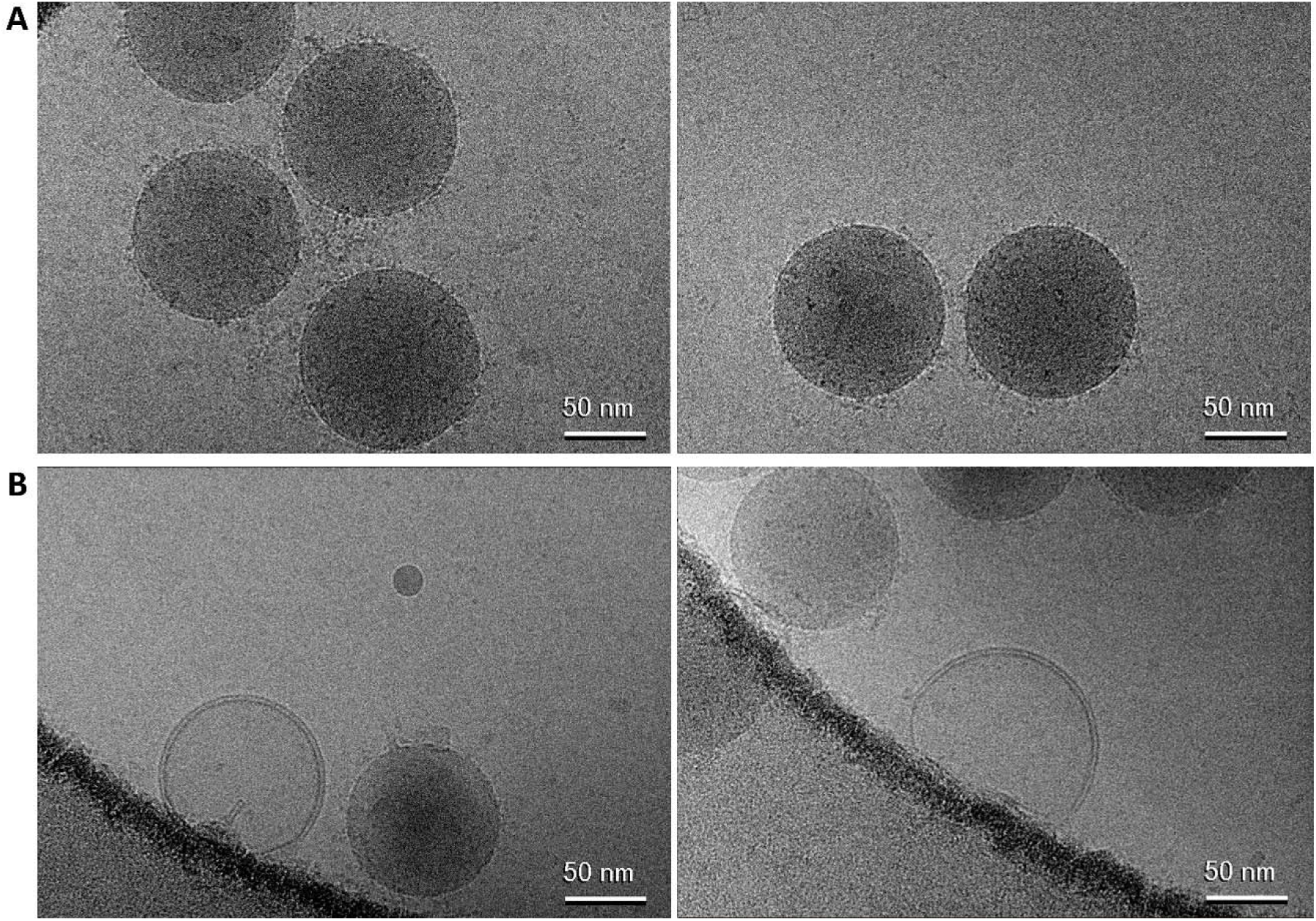
Competitive comparison of divergent protein corona architectures on hard *vs.* soft NPs. **A)** Representative cryo-TEM images of solid-core polystyrene NPs incubated with purified ApoE4. The images show the formation of a robust, classical “fuzzy” protein corona shell (characteristic peripheral electron density) completely enveloping the hard particle surfaces. **B)** Representative cryo-TEM images where polystyrene NPs and DOTAP LNPs were mixed in the same vessel and simultaneously exposed to purified ApoE4. Crucially, within the same field of view, the polystyrene NPs develop a traditional dense protein corona shell, whereas the neighboring LNPs lack any detectable fuzzy shell. Instead, the LNPs show evidence of distinct localized membrane thickening and ApoE4-bilayer integration.

We then conducted a “dual-particle” experiment by mixing polystyrene NPs and DOTAP LNPs in a single volume and incubating the mixture with purified ApoE4, ensuring that both particle types are exposed to identical ApoE4 concentrations and environmental conditions. As shown in **Fig. 4B**, our results revealed a striking contrast in protein organization within the same field of view: while the solid-core PSNPs developed a distinct, dense protein shell, the neighboring hollow-core LNPs entirely lacked this architecture. Instead, the LNPs exhibited localized membrane thickening and electron-dense integration sites, indicating that even in a direct competitive environment, ApoE4 engages the LNP through localized membrane remodeling rather than shell formation. This side-by-side comparison supports a model in which the LNP surface is remodeled rather than coated.

To further investigate the molecular basis of ApoE4-DOTAP LNP interactions, we performed all-atom molecular dynamics (MD) simulations of two ApoE4 constructs in the presence of a DOTAP lipid bilayer (**Fig. S1A**). The first construct, the 22 kDa N-terminal domain (NTD, residues 22-165, PDB ID: 1GS9), was used to characterize how the receptor-binding four-helix bundle engages the bilayer in isolation. The second construct was the full-length (FL) ApoE4 (residues 1-299), generated by reverting the five solubilizing mutations (F257A, W264R, V269A, L279Q, V287E) in the monomeric ApoE3 NMR structure (PDB ID: 2L7B) and introducing the C112R substitution to convert the sequence to the ApoE4 isoform.

For each construct, three protein-bilayer systems were prepared with distinct initial orientations to sample different engagement pathways (**Fig. S1B**): (i) the OPM/PPM-predicted membrane-binding orientation; (ii) the same orientation translated 20 Å above the bilayer to test spontaneous association from the aqueous phase; and (iii) the OPM/PPM orientation rotated 180° about the y-axis to invert the protein. Three additional control simulations (NTD-only, FL-only, and DOTAP-only) provided baselines for protein conformational dynamics and unperturbed bilayer properties.

System equilibration was assessed by root mean square deviation (RMSD) and root mean square fluctuation (RMSF) of the protein backbone (**Fig. S2**). The NTD construct exhibited stable backbone RMSD values of ∼2 Å across all systems, consistent with the rigidity of its four-helix bundle (**Fig. S2A**). Per-residue Cα RMSF showed correspondingly low fluctuations in the helical segments and higher fluctuations in the inter-helical loops, with similar patterns across the three replicates (**Fig. S2C**). In contrast, the FL construct showed substantially larger RMSD values (**Fig. S2B**), reflecting reorganization of its intrinsically disordered regions (the hinge, residues 166-243, and the C-terminal tail, residues 278-299). The RMSF analysis confirms this interpretation, localizing the dynamics to the disordered hinge and C-terminal tail regions while the structured four-helix bundle remained stable across all replicates. The amphipathic helix also remained stable in Rep_B and Rep_C, but showed elevated fluctuations in Rep_A (**Fig. S2D**)

Across all replicates, the NTD established sustained contacts with the bilayer, with the minimum protein-lipid distance settling at ∼2 Å throughout the trajectories (**Fig. S3A**). Per-residue contact frequency analysis identified the H1-H2 face as the consistent binding interface across replicates (**Fig. S3B**, **Fig. 5A**). Decomposing these contacts by residue type revealed that acidic and aromatic residues bind most frequently on a per-residue basis, while acidic and polar residues contribute the largest total interaction surface (**Fig. S3C**).

**Fig. 5.**
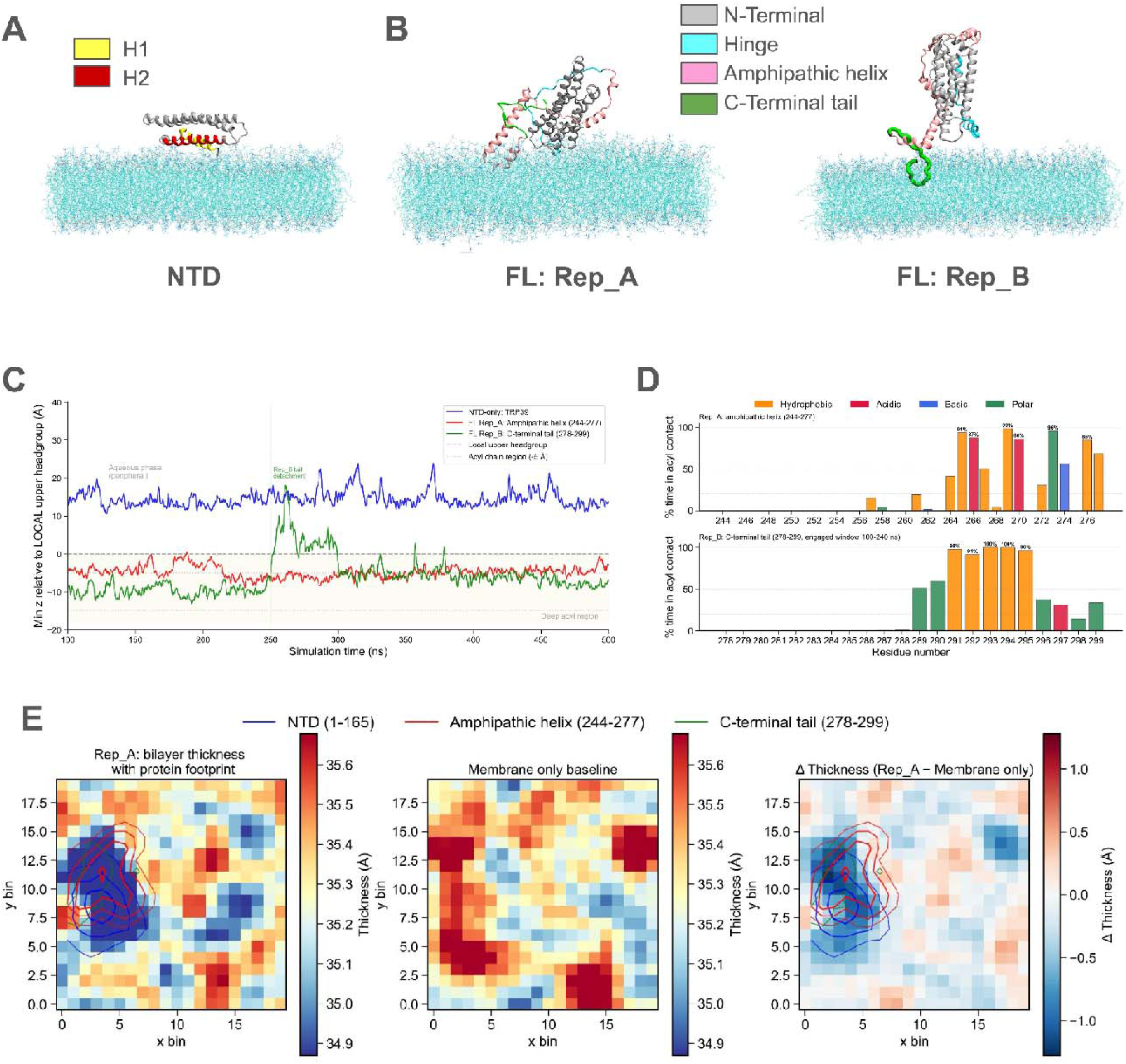
Molecular dynamics analysis of ApoE4-DOTAP bilayer interactions reveals orientation-dependent partial embedding via the C-terminal domain. **A)** Late-frame snapshot of the N-terminal domain (NTD) showing peripheral engagement at the bilayer surface via the H1-H2 face (H1 yellow, H2 red); no insertion observed. **B)** Late-frame snapshots of two full-length ApoE4 (FL) replicates showing distinct insertion modes. Rep_A (left): partial embedding of the amphipathic helix (244-277, pink). Rep_B (right): deep insertion of the C-terminal tail (278-299, green). NTD shown in gray, hinge in cyan. **C)** Time-resolved minimum z-position relative to the local upper headgroup plane for NTD-TRP39 (blue), Rep_A amphipathic helix (red), and Rep_B C-terminal tail (green), showing peripheral, intermediate, and deep insertion respectively. The tail dissociates from Rep_B at ∼250 ns. **D)** Per-residue percentage of frames in acyl chain contact (<4 Å) for the Rep_A amphipathic helix (top) and Rep_B C-terminal tail (bottom, engaged window 100-240 ns), color-coded by residue type: hydrophobic (orange), acidic (red), basic (blue), polar (green). Helix anchors include aromatic (PHE265), acidic (GLU266), and aliphatic (VAL269) residues; tail anchors are purely aliphatic (ALA291, ALA292, PRO293, VAL294, PRO295). **E)** Bilayer thickness perturbation for Rep_A. Left: thickness with protein engagement. Center: membrane-only baseline. Right: Δ thickness showing localized thinning (∼0.5 Å) beneath the helix engagement site. Protein contact footprints overlaid as contours: NTD (blue), amphipathic helix (red), C-terminal tail (green).

To characterize the underlying interactions, we computed the total number of hydrogen bonds between NTD and DOTAP and found this to be modest across all replicates (**Fig. S4A**). We therefore examined the top contact residues individually (**Fig. S4B**), identifying three glutamines (GLN46, GLN55, GLN81), three glutamates (GLU49, GLU50, GLU80), and a single tryptophan (TRP39) as the dominant binding residues. The GLN residues showed limited direct hydrogen bonding with DOTAP (**Fig. S4C**). The GLU residues engaged DOTAP predominantly through water-mediated electrostatic interactions rather than direct salt bridges (**Fig. S4D**). TRP39 consistently formed cation-π interactions with DOTAP trimethylammonium headgroups across all three replicates (**Fig. S4E**). Together, these analyses indicate that NTD engages the bilayer through a network of hydrogen bonds, water-mediated electrostatics, and cation-π interactions, anchored at the H1-H2 face. Consistent with this peripheral engagement, bilayer thickness analysis showed only modest local thinning without disruption of bilayer integrity (**Fig. S5**).

Contact analysis between FL ApoE4 and DOTAP revealed persistent protein-bilayer interactions across all three replicates, with only a brief detachment event in Rep_B at ∼250 ns (**Fig. S6A**). Per-residue contact frequency analysis identified three distinct orientation-dependent binding modes (**Fig. S6B**). Rep_A engaged the bilayer primarily through residues in the CTD lipid-binding region (244-277), consistent with the elevated RMSF observed for this region in this replicate. Rep_B engaged primarily through the C-terminal tail (278-299), while Rep_C showed dispersed contacts across both N- and C-terminal regions. Decomposing these contacts by residue type identified aromatic and aliphatic hydrophobic residues as the dominant contributors across all three simulations (**Fig. S7**).

To quantify the engagement geometry of each binding mode, we computed the minimum z-position of the amphipathic helix and C-terminal tail relative to the local upper headgroup plane (**Fig. S8**, **Fig. 5C**). In Rep_A, the amphipathic helix was continuously inserted at the bilayer interface while the C-terminal tail remained in the aqueous phase. Rep_B showed the inverse pattern, with the C-terminal tail penetrating deeply into the bilayer while the amphipathic helix remained above, until the tail detached at ∼250 ns. The protein subsequently re-engaged in a sparser configuration, where the minimum z-position recovered to comparable values but only a small subset of residues from each region contacted the bilayer (trajectory inspection, **Fig. 5B**). Rep_C showed both regions remaining above the bilayer throughout the simulation, consistent with the inverted starting orientation preventing sustained CTD engagement.

We also computed per-residue local-reference depth profiles for the amphipathic helix and C-terminal tail regions, to identify the specific residues anchoring each engagement mode (**Fig. S9**). In Rep_A, residues PHE265, GLU266, VAL269, GLN273, and TRP276 of the amphipathic helix were consistently positioned at or below the local headgroup plane, while the C-terminal tail residues remained above. In Rep_B, C-terminal tail residues 290-296 sustained deep insertion into the upper acyl region, with the hydrophobic cluster at residues 291-295 (ALA, ALA, PRO, VAL, PRO) achieving the deepest penetration. Per-residue acyl chain contact analysis (**Fig. S10**, **Fig. 5D**) confirmed these anchoring residues, with sustained acyl contact and multiple heavy atoms simultaneously embedded in the hydrocarbon region.

Bilayer thickness analysis revealed replicate-specific membrane perturbations correlating with each engagement mode (**Fig. S11**). Rep_A showed localized bilayer thinning beneath the amphipathic helix footprint, consistent with surface-aligned helix engagement compressing the upper leaflet. Rep_B showed localized thickening beneath the C-terminal tail footprint, consistent with deep tail insertion displacing headgroups outward. Rep_C showed modest thickening under the NTD-down footprint. Together, these contrasting bilayer signatures reflect the distinct molecular geometries of the three engagement modes and provide a molecular basis for the heterogeneous bilayer remodeling observed by cryo-TEM.

While the bilayer perturbations observed in our simulations are modest and full integration of ApoE4 into the bilayer was not achieved within the accessible timescales, our results nonetheless support the partial embedding mechanism inferred from the cryo-TEM measurements. Complete protein integration into the bilayer, as depicted in our proposed mechanism (**Fig. 2F**), likely requires biological timescales beyond those accessible to atomistic MD simulations. Additional factors not captured by our simulation setup, including cooperative effects of multiple ApoE4 proteins binding to the same LNP, and the high local curvature characteristic of LNP surfaces, may further amplify these perturbations and facilitate the full integration observed experimentally.

Lipoproteins and EVs are known to form functional complexes in various biofluids.^27, 28^ Interactions between multiple types of (apo)lipoproteins such as ApoE (found in chylomicrons, high-density lipoprotein (HDL), and VLDL), ApoM (in HDL), and ApoCII and ApoCI (in chylomicrons, HDL, and VLDL), with lipid-based nanoparticles including EVs, LNPs, Doxil, and LIPO are well-documented.^29^ This fusion, along with the potential protein interactions with blebs, likely contributes to the efficient uptake of LNPs by liver hepatocytes, mediated by the interaction of ApoE-rich lipoproteins with the nanoparticle surface following intravenous administration.^27^

Many proteins associated with LNPs likely arise from specific lipid-driven interactions, either through lipid bilayer fusion or interaction with blebs, as opposed to protein adsorption observed with hard nanoparticles. It has been previously established that lipoprotein components can interact with plasma proteins through hydrophobic insertion.^28^ Lipidated proteins bearing palmitoyl, myristoyl, or prenyl groups, insert hydrophobically into membranes; palmitoylation functions as a reversible tether that regulates trafficking, whereas prenylation provides a largely irreversible anchor.^30, 31^ Studies show farnesylated peptides and lipidated motifs spontaneously insert into bilayers and exhibit rapid intermembrane exchange with domain preferences dictated by lipid composition.^32–34^ Because the outer leaflet of LNPs is rich in phospholipids, cholesterol, and PEG-lipids, similar hydrophobic principles enable lipidated proteins or peptides to embed in, or associate with, LNP surfaces. Consistent with the latter mechanism, ApoE adsorbs to circulating LNPs, reorganizes LNP architecture, and mediates hepatocyte uptake via LDL receptors, a key determinant for siRNA/mRNA delivery.^29^

Our structural findings provide mechanistic insight into recent functional studies of LNP protein coronas. Previous studies have demonstrated that plasma proteins, particularly apolipoproteins such as ApoE, enrich on LNP surfaces after plasma exposure and significantly influence LNP cellular uptake, intracellular trafficking, and mRNA transfection efficiency.^35^ For instance, Voke *et al.*^36^ recently showed that three of the proteins identified by our MS analysis - vitronectin, C-reactive protein, and alpha-2-macroglobulin - associate with LNPs and modulate delivery outcomes. Interestingly, the study also demonstrated that some proteins increase uptake while paradoxically decreasing transfection, possibly through altered lysosomal trafficking.

Our cryo-TEM data offer a structural basis for these observations: rather than forming discrete peripheral shells that might act as barriers, proteins appear to associate with LNPs through lipoprotein fusion and membrane integration. This fusion-based mechanism has several functional implications. First, lipoprotein fusion directly alters LNP membrane composition and physical properties, potentially affecting membrane fluidity, stability, and fusogenicity with endosomal membranes. This could explain why protein association affects not only uptake but also downstream events like endosomal escape. Second, apolipoprotein-enriched LNPs are recognized by lipoprotein receptors (e.g., LDL receptors for ApoE-bearing particles), providing a molecular mechanism for the liver tropism commonly observed with intravenously administered LNPs.^35^ Our observation that lipoprotein fusion occurs consistently across LNP formulations supports this as a general mechanism of protein-mediated targeting. Third, the dynamic nature of membrane fusion, as opposed to static shell formation, may allow for protein exchange and remodeling during circulation. This could potentially explain the evolution of corona composition observed in kinetic studies. The fusion mechanism also suggests opportunities for rational design: rather than preventing protein adsorption (the typical goal with PEGylation), LNP formulations could be optimized to favor specific lipoprotein interactions that enhance targeting to desired tissues.

Our findings revise the structural picture of how proteins interact with LNPs, from a conventional adsorbed shell toward lipoprotein fusion, association, and potential bleb interactions. This shift affects biodistribution: to achieve cell-specific targeting, we must recognize lipoprotein-mediated uptake (e.g., via ApoE/LDL receptor binding) as a primary mechanism. This then necessitates the engineering of specific lipoprotein coronae for targeted delivery, achievable by modulating LNP lipid composition or incorporating ligands like ApoE-mimetics. Our mechanistic insights reframe existing design principles: rather than empirically balancing PEGylation against cellular uptake, formulations can now be rationally designed to modulate specific lipoprotein fusion events through coordinated optimization of PEG content, lipid composition (helper, ionizable lipids), and surface charge. Finally, it influences stability and cargo release: the intimate nature of lipoprotein fusion affects LNP stability, membrane integrity, and encapsulated cargo release, making it crucial to design LNPs that leverage productive fusion for endosomal escape or payload delivery, and to develop new *in vitro* and *in vivo* assays to optimize these specific fusion and bleb interactions.

In summary, our results challenge the prevailing assumption that soft, lipid-based nanoparticles, such as LNPs, acquire a traditional protein corona, which has been observed as a discrete, adsorbed protein shell along the surface of harder nanoparticles, such as polystyrene nanoparticles. Instead, our structural and computational data support a model in which LNPs engage plasma proteins through lipoprotein fusion or interactions with bleb structures, forming hybrid assemblies in which lipoproteins integrate into the LNP surface. Whereas a traditional protein corona passively coats the nanoparticle surface and may desorb or reorganize dynamically, this lipoprotein fusion implies a stable, compositional transformation of the lipid-based nanoparticles. Consequently, the biological identity of the LNP may not be merely masked by serum components but substantially altered by the incorporation of endogenous lipoproteins. This can effectively embed LNPs within endogenous lipid transport processes in vivo, enabling engagement with native lipid trafficking and receptor-mediated uptake pathways while potentially reducing immune recognition. It is noteworthy that these mechanistic findings are based on one ionizable lipid, and generalization to two-tailed lipids like DLin-MC3-DMA will require further analysis.

This shift reframes how LNPs interact with biological systems, moving from generic protein adsorption to specific lipoprotein fusion events that can be rationally engineered. These findings necessitate an expanded view of biological interactions for lipid-based nanomaterials, in which biological identity may emerge through membrane remodeling rather than shell-like adsorption. For LNPs, formulation parameters such as ionizable lipid chemistry, cholesterol content, and helper lipid selection are likely to dictate the extent and specificity of lipoprotein fusion, rather than minimizing plasma interactions, LNP formulations may be deliberately engineered to exploit selective fusion with specific lipoprotein classes, thereby tuning biodistribution, cellular uptake, and circulation stability. To provide further data on how to advance rational lipid-based nanomedicine design, future studies should 1) characterize fusion kinetics and composition for different lipoprotein classes; 2) correlate specific lipoprotein interactions with in vivo biodistribution; 3) develop structure-function relationships between lipid composition and lipoprotein fusion; 4) explore how disease-specific changes in plasma lipoprotein profiles^13, 28^ affect LNP protein corona composition and targeting; and 5) engineer LNP formulations that actively recruit therapeutically beneficial lipoproteins. Further work should also focus on decoding the exact lipoprotein motifs that drive lipid membrane fusion, as these motifs may be conserved within glycoproteins, such as vitronectin and beta 2 glycoprotein I (β2-GPI), which have been shown to mediate LNP trafficking to the lungs and spleen, respectively.^37^ Related glycoprotein-mediated targeting has also been reported for the placenta. ^38^ Once the protein-lipid interactions that drive plasma proteins and LNP fusion are decoded, perhaps new ionizable lipids or helper lipids could be designed that encourage or directly mediate the fusion of precise, organ-targeting lipoproteins within LNPs upon systemic administration.

Overall, recognizing lipid-based nanoparticles as dynamic, biologically integrative assemblies will be critical for improving *in vivo* biodistribution predictability and provides a new framework for engineering next-generation targeted nanomedicines across a broad range of therapeutic applications.

## Materials and Methods

The experimental methods for the preparation of the protein corona, as well as the mass spectrometry analysis of both the EVs and the protein corona, along with the electron microscopy analysis of the LNPs and molecular dynamic simulation, are detailed in the **SI**.

## Appendix

### Data Availability

Raw mass spectrometry data: <u>PRIDE</u>, Project accession PXD070622 (Token: AduYuUTzOIXz). Processed proteomics data are available in Supplementary Data Sets 1 (for EVs) and 2 (for LNPs).

### Contributions statement

S.G., J.C., S.A.M.S., F.F., K.H., and B.S. developed workflows and performed experiments for proteinl71corona formation, isolation, and characterization of coronal71coated LNPs. N.M. synthesized and characterized LNPs. G.Y. and M.R.K.M. performed molecular dynamics analyses. A.A.S., A.M.M., M.J.M., B.B., and L.S. contributed to discussions of the mechanistic findings and their implications for the field. H.V. performed cryol71TEM imaging and analysis. K.A.W. planned and supervised LNP formulation and contributed to interpretation and revision. M.M. conceived the study, planned experiments, and supervised proteinl71corona work. All authors contributed to writing and revising the manuscript and approved the final version.

### Supporting Information

Detailed experimental protocols for LNP synthesis, protein corona formation and characterization, mass spectrometry-based proteomic analysis, and molecular dynamics.

