## Supplementary material for "Lipid nanoparticle protein coronas arise through lipoprotein fusion rather than shell-like adsorption": SI

**Materials and methods**

**LNP formulation.** Ionizable lipid nanoparticles (LNPs) were formulated as previously described ^1^. Briefly, the ionizable lipid 306O_i10_, helper phospholipid, cholesterol, and C14-PEG2000 were mixed at a standard molar ratio of 35:16:46.5:2.5 and then dissolved in ethanol and 10 mM sodium citrate. Four different helper lipids – DOPE, DSPC, DOTAP, and DOPS – were used to create nanoparticles. For DOPE-containing nanoparticles, the two additional molar ratios - 35:16:47.5:1.5 and 35:16:49:0 – were tested. mRNA was dissolved in 10 mM sodium citrate. The lipid solution and mRNA were rapidly mixed via pipetting and the final lipid:mRNA ratio by weight was 10:1. All nanoparticles were dialyzed in phosphate-buffered saline for 90 minutes using 3500 MWCO dialysis cassettes (Thermo Fisher). The characteristics of the synthesized LNPs are summarized in T**able S1**.

**Table S1.** Characteristics of the synthesized DOPE, DSPC, DOTAP, and DOPS LNPs.

| Characterization | DOPE LNP | DOTAP LNP | DSPC LNP | DOPS LNP |
| --- | --- | --- | --- | --- |
| Average size | 103 nm | 100 nm | 85 nm | 95 nm |
| Average Zeta Potential | -4.02 | -7.71 | -6.19 | -10.3 |
| Average PDI | 0.28 | 0.35 | 0.11 | 0.24 |
| Average RNA Entrapment | 90.4% | 82.4% | 36.8% | 24.7% |
| Average pKa | 6.3 | 6.4 | 6.2 | 6.3 |

Size, zeta potential, and PDI were measured by a Malvern Zetasizer. RNA entrapment was measured using a Quant-iT™ RiboGreen™ RNA Assay Kit according to manufacturer instructions. Surface charge, or pKa, was measured using a Sodium 2-(p-toluidino)-6-naphthalenesulfonic acid (TNS) ionization assay.

**Polystyrene nanoparticles.** Two different sizes (i.e., 100 nm and 200 nm) of carboxylated polystyrene NPs were purchased from Polysciences ([www.polysciences.com](http://www.polysciences.com)).

**Human plasma.** Pooled healthy human plasma was purchased from BioIVT Company ([www.bioivt.com](http://www.bioivt.com)).

Apolipoprotein E (ApoE). Recombinant human-derived APOE protein, expressed by HEK293, was purchased from MedChemExpress (HY-P78542).

**EV separation.** A fixed volume of exosome sample (20uL) was used for MACSPlex analysis using MACSPlex EV Kit, enabling the characterization of 37 EV surface markers in a single experiment. The screening approach is based on fluorescent immune conjugates—MACSPlex EV Capture Beads—which bind different EV-specific epitopes. Kit’s reagent marks the bound exosomes, for example, with the pan-exosome markers CD9, CD63 and CD81. The result is a sandwich complex, consisting of the MACSPlex Exosome Capture Bead, the exosome and the detection reagent, which can subsequently be analyzed based on the respective fluorescent properties. Using flow cytometry, semiquantitative data on 37 EV surface epitopes are obtained. More specifically, the flow cytometer was calibrated, and background settings were adjusted to unlabeled beads and gating strategies were used to identify bead populations for each analyte. Batch analysis quantified median intensities for each bead population and analyte surface expression was calculated for each sample.

**Proteomics analysis of the collected EVs.**

*Lysing and digestion of the EV proteins.* EVs were lysed using a buffer containing 50 mM Tris, 100 mM NaCl, 2% SDS, 10 mM EDTA, and 40 mM water-based solution. The EVs (1:2 volume ratio) were incubated and shaken at 60°C for 60 minutes to disrupt cell membranes. Followed by alkylation, 25 mM iodoacetamide was added and the sample was incubated in the dark at room temperature for 30 minutes. The lysate was loaded onto a 5 kDa cutoff filter and centrifuged to remove the lysis buffer, then washed twice with 25 mM Tris and 100 mM NaCl. The final volume was adjusted to 100 μL with the same buffer. Proteins were digested on the filter with 20 μg of trypsin-LysC overnight at 38°C. Reaction was stopped by adding 5 μL of formic acid, and the supernatant containing peptides was collected after a 30-minute centrifugation.

*LC/MS/MS analysis of digested proteins.* Peptides were separated on an Agilent 2.1×100 mm, 2.7 μm C18 column using a linear gradient of 0–50% B (A = water + 0.025% TFA; B = 95% acetonitrile + 0.025% TFA) over 60 minutes at 0.3 mL/min. The column was held at 40°C with a 30 μL injection volume. Analysis was performed on an Agilent 6530 q-TOF MS in positive mode, covering m/z 300–3000 in full MS scans. MS/MS spectra were acquired for the top two precursor ions (above 8000 counts, m/z 500–2000) using nitrogen collision gas, with collision energies calculated as:

CE= (3.6 **m/z* /100)-4.8

Instrument parameters included a capillary voltage of 3700 V, nozzle at 1500 V, fragmentor at 160 V, drying gas at 350°C (8 L/min), nebulizer at 40 psi, and sheath gas at 350°C (12 L/min). The first 3 minutes of each run were diverted to waste. Reference masses at m/z 322, 922, and 1522 ensured calibration, delivered via a secondary nebulizer with a 0.5 mL/min flow split. Data acquisition used Agilent MassHunter software, and protein identification was performed with Spectrum Mill (Rev B.04.00.127) against the UniProtKB/Swiss-Prot human proteome database, with BSA also searched against mammalian proteins.

**Protein corona sample preparation for LNPs.** To collect LNPs after plasma interaction, we used size exclusion chromatography with a Sepharose CL-4B column (10×1.5 cm). For corona formation, 450 μL of stock LNPs (1 mg/mL) was mixed with 550 μL of human plasma and incubated at 37°C for 1 hour with shaking in a thermomixer for uniform coating. The mixture was then diluted to 1 mL with PBS and loaded onto the column. After collecting 3 mL containing the corona-coated LNPs, an additional 15 mL of PBS was passed through the column to elute remaining plasma proteins. To determine when plasma proteins elute, plasma alone was loaded, fractions collected, and analyzed by gel electrophoresis. Results showed plasma proteins begin to elute from fraction 7 onwards. Based on this and supported by DLS analysis (and in line with other reports ^2^), we collected early, plasma-free fractions (below fraction 7), as larger LNP particles pass through the column faster than smaller plasma proteins. The purified LNPs were further cleaned of loosely attached proteins using a Vivaspin 500 concentrator (MWCO 1,000,000). Centrifugation retains the LNPs at the top, while free or loosely bound plasma proteins are removed at the bottom. The collected LNPs then underwent peptide preparation for mass spectrometry analysis.

**Peptide preparation for mass spectrometry.** The collected LNPs were resuspended in 20 µL of PBS containing 0.5 M guanidinium hydrochloride. Proteins were reduced with 2 µL of 2 mM DTT (from 1 M stock) and incubated at 37°C for 45 minutes, followed by alkylation with 2.2 µL of 8 mM IAA (from 1 M stock) for 45 minutes at room temperature in the dark. Afterward, 5 µL of LysC (0.02 µg/µL in PBS) was added, and the mixture was incubated for 4 hours at 37°C. This was followed by overnight digestion with trypsin at 37°C at the same concentration. The samples were then centrifuged at 12,000 g for 5 minutes at room temperature using Vivaspin 500 filter columns, collecting the bottom fraction to remove the LNPs. The peptides were acidified to pH 2-3 with TFA and purified using C18 cartridges.

**Protein corona sample preparation for polystyrene nanoparticles.** Buffer-diluted polystyrene nanoparticles (0.62 mg/mL) were mixed with 55% human plasma. The nanoparticle-plasma mixtures (total volume: 500 μL) were incubated at 37°C for 1 hour with constant agitation at 10 G. After incubation, the samples were centrifuged at 2000 × g for 30 minutes at 10–15°C to pellet the polystyrene nanoparticle–protein complexes. To remove the soft corona, the pellets were resuspended in 400 μL of cold Sorensen’s phosphate buffer and centrifuged again for 10 minutes. To minimize contamination from low-affinity proteins, the pellets were washed three times and finally resuspended in 200 μL of Sorensen’s phosphate buffer for cryo-TEM analysis.

**Preparation of ApoE protein corona.** Aliquots (100 µL) of DOTAP LNPs or 100 nm polystyrene nanoparticles (1 mg/mL) were mixed with 50 µL of ApoE and incubated for 1 hour at 37°C with agitation. The samples were then processed for cryo-TEM imaging. To assess variations in corona formation across different particles, both DOTAP LNPs and polystyrene nanoparticles were also mixed together prior to the addition of ApoE.

**Mass spectrometry data processing of protein corona.** Bottom-up MS data were analyzed using MaxQuant (v1.5.5.1) for peptide and protein identification with label-free quantification (LFQ) based on database searching. Using MaxQuant, MS raw files were analyzed LFQ enabled. Data were searched against the UniProt human proteome database (UP000005640; 82,733 entries; version dated December 29, 2023). A decoy (reversed) database was used to estimate false discovery rates (FDR). The search parameters specified trypsin as the proteolytic enzyme, allowing up to two missed cleavages. Carbamidomethylation of cysteine residues was set as a fixed modification, while methionine oxidation, deamidation of asparagine/glutamine, and N-terminal acetylation were treated as variable modifications. Filtering criteria included a 1% FDR threshold at both the peptide-spectrum match (PSM) and protein group levels. All other parameters followed the software’s default settings. Subsequent data analysis was conducted using Perseus software (v2.0.10.0). Entries annotated as "Reverse," "Potential contaminant," or "Only identified by site" were excluded from downstream analysis. Heatmaps was generated using an S₀ value of 0.05 and an FDR of 0.1.

**Dynamic light scattering (DLS).** DLS analysis was performed to determine the size distribution of the LNPs before incubation with plasma and after collecting following plasma exposure, using a Zetasizer Nano Series instrument (Malvern). Measurements were conducted at room temperature with a 632 nm Helium-Neon laser.

**Gel electrophoresis.** Sodium dodecyl sulfate–polyacrylamide gel electrophoresis (SDS-PAGE) was performed on the fractions collected from the column to determine where plasma proteins begin eluting. For each collected fraction, 20 μL was mixed with 20 μL of 2× Laemmli sample buffer, heated at 85°C for 6 minutes, and loaded onto precast gels. After electrophoresis, the gels were fixed in a solution containing 10% acetic acid and 40% ethanol, then stained overnight with 50 mL of Coomassie blue stain. Following multiple washes, the gels were scanned the next morning for analysis.

**Cryo-TEM.** 10-nm BSA-treated gold nanoparticles were mixed with each of the LNPs and polystyrene nanoparticles at a ratio of 1:4.3. Five microliters of each sample were applied to glow-discharged holey carbon grids (C-Flat R2/2, Protochips, Inc.), blotted, and rapidly frozen in liquid ethane using the Vitrobot Mark IV (Thermo Fisher Scientific, Hillsboro, OR, USA). Images were acquired on a Titan Krios 300 kV Cryo-Transmission Electron Microscope equipped with a Falcon 2 direct electron detector and phase plate (Thermo Fisher Scientific). The microscope operated at a nominal magnification of 75,000×, resulting in a pixel size of 1.075 Å, with defocus values ranging from −2.0 to −3.0 μm under low-dose conditions.

**Molecular dynamics simulation components and setups.** Two ApoE4 protein systems were prepared. For the N-terminal domain (NTD) simulations, the crystal structure of the 22 kDa NTD fragment (PDB ID: 1GS9, residues 22-165)^3^ was used directly. For the full-length ApoE4 (FL, residues 1-299) simulations, the NMR structure of a monomeric ApoE3 variant (PDB ID: 2L7B, model 1)^4^ was used as a template. This monomeric variant contains five engineered mutations (F257A, W264R, V269A, L279Q, V287E) designed to prevent oligomerization. These mutations were reversed to recover the wild-type ApoE3 sequence, after which the C112R substitution was introduced to convert the sequence to ApoE4 (Fig. S1A).

Symmetric DOTAP bilayers (14 × 14 nm in the xy plane) were constructed using the CHARMM-GUI Membrane Builder ^5–9^. Three protein-bilayer systems were prepared for each protein with distinct initial protein orientations relative to the bilayer (Fig. S1B): Rep_A used the orientation predicted by the OPM/PPM server^10^, with the protein positioned at the predicted membrane-binding interface; Rep_B used the same PPM orientation translated +20 Å along the bilayer normal, placing the protein in the aqueous phase above the bilayer; Rep_C used the PPM orientation rotated 180° about the y-axis, inverting the protein orientation relative to the bilayer. These distinct initial conditions were designed to sample multiple protein-bilayer encounter pathways.

Three additional control simulations were performed for baseline comparisons: an NTD-only simulation (NTD solvated without bilayer), an FL-only simulation (FL solvated without bilayer), and a membrane-only DOTAP bilayer simulation (without protein).

**Molecular dynamics simulation protocol.** All simulations were performed using GROMACS^11^ with the CHARMM36m^12^ force field. Systems were solvated with the TIP3P water model and neutralized with 0.15 M NaCl. Hydrogen mass repartitioning^13^ (HMR) was applied during CHARMM-GUI system construction to enable a 4 fs production time step.

Energy minimization was performed using the steepest descent algorithm for 5000 steps or until the maximum force fell below 1000 kJ/mol/nm. Equilibration of the protein-bilayer and membrane-only systems followed the standard CHARMM-GUI six-step protocol: three NVT equilibration steps (125 ps each, 1 fs time step) followed by three NPT equilibration steps (500 ps each, 2 fs time step), during which position restraints on protein backbone atoms, protein side chains, and lipid headgroups were progressively reduced. The protein-only control systems underwent a simpler equilibration consisting of one NVT step and one NPT step (125 ps each, 1 fs time step).

Production simulations were run for 500 ns per system at 310 K and 1 atm with a 4 fs time step. Temperature was maintained using the v-rescale thermostat (τ = 1.0 ps) with separate coupling groups for protein, lipids, and solvent. Pressure was maintained semi-isotropically using the c-rescale barostat (τ = 5.0 ps, compressibility = 4.5 × 10⁻⁵ bar⁻¹). Long-range electrostatics were treated with the particle mesh Ewald (PME) method with a real-space cutoff of 1.2 nm. Van der Waals interactions were truncated at 1.2 nm using a force-switch function starting at 1.0 nm. Bonds involving hydrogen atoms were constrained using LINCS. Periodic boundary conditions were applied in all three dimensions.

The first 100 ns of each production simulation were treated as additional unrestrained equilibration and discarded from analysis. All analyses were performed on the last 400 ns of the trajectories unless stated otherwise.

**Molecular dynamics post-processing analysis.** Standard trajectory analyses (RMSD, RMSF, minimum distance, number of H-bonds) were performed using GROMACS analysis tools. All custom analyses (contact frequency, depth profiles, hydrophobic contacts, bilayer thickness perturbation) were implemented in Python using MDAnalysis^14^ and LiPyphilic^15^ . The first 100 ns of each simulation were discarded as additional unrestrained equilibration; all subsequent analyses were performed on the remaining production trajectories.

Backbone RMSD and per-residue Cα RMSF were computed using gmx rms and gmx rmsf after least-squares fitting to the first frame of the production. The minimum distance between protein and DOTAP heavy atoms was computed using gmx mindist.

Hydrogen bonds between protein and DOTAP atoms were computed using gmx hbond with default geometric criteria (donor-acceptor distance < 3.5 Å, angle cutoff 30° from linearity).

Contact frequency was computed per residue as the fraction of frames in which any residue heavy atom was within 6.0 Å of any DOTAP heavy atom.

Per-residue depth was computed relative to the local upper headgroup, defined per atom as the median z of the 5 nearest upper DOTAP nitrogen atoms in the xy plane. Insertion was quantified as the fraction of frames in which the residue's deepest atom was below this local reference. For the time-resolved analyses (Fig. 5C), the minimum depth across all atoms in the specified region (NTD TRP39 ring, amphipathic helix, or C-terminal tail) was reported per frame.

Hydrophobic contact was computed as the fraction of frames in which any residue heavy atom was within 4.0 Å of a DOTAP acyl chain carbon (C22-C218 and C32-C318), providing direct evidence of contact with the bilayer hydrocarbon region.

Bilayer thickness was computed using LiPyphilic's AssignLeaflets and MembThickness on a 20 × 20 grid with linear interpolation. Δ thickness maps were computed as the difference between protein-bilayer and matched-window membrane-only simulations.

For FL Rep_B, analyses involving the C-terminal tail engagement (per-residue depth, hydrophobic contact, bilayer thickness) were restricted to the engaged window (100-240 ns), prior to the tail dissociation at ~250 ns.

All MD visualizations were done with VMD^16^ and plots made with python.

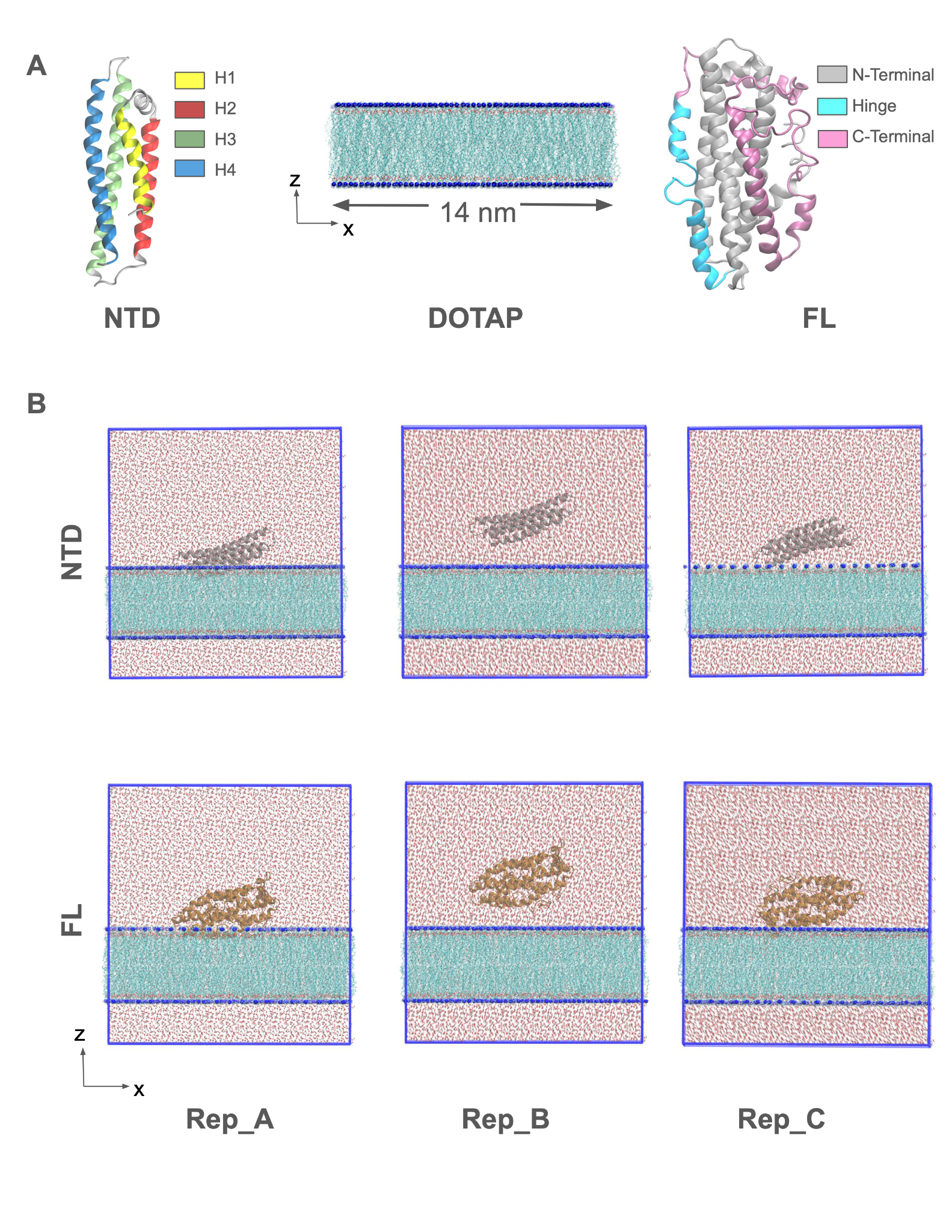

**Fig S1. Molecular dynamics simulation system setup. A)** Schematic representation of the simulation components, including the four-helix bundle of the ApoE4 N-terminal domain (NTD), the DOTAP bilayer, and full-length ApoE4 (FL). Helices H1–H4 of the NTD are highlighted and color-coded. For FL ApoE4, the N-terminal domain, hinge region, and C-terminal domain are indicated. **B)** Schematic of the initial protein–membrane simulation configurations. The top row corresponds to the NTD systems and the bottom row to the FL systems. From left to right, the starting conformations include: (i) the orientation predicted by the PPM server (Rep_A), (ii) the same orientation translated 20 Å away from the membrane surface (Rep_B), and (iii) the PPM orientation rotated 180° about the y-axis (Rep_C). In addition to these simulations, a protein only system and a membrane only system were also run.

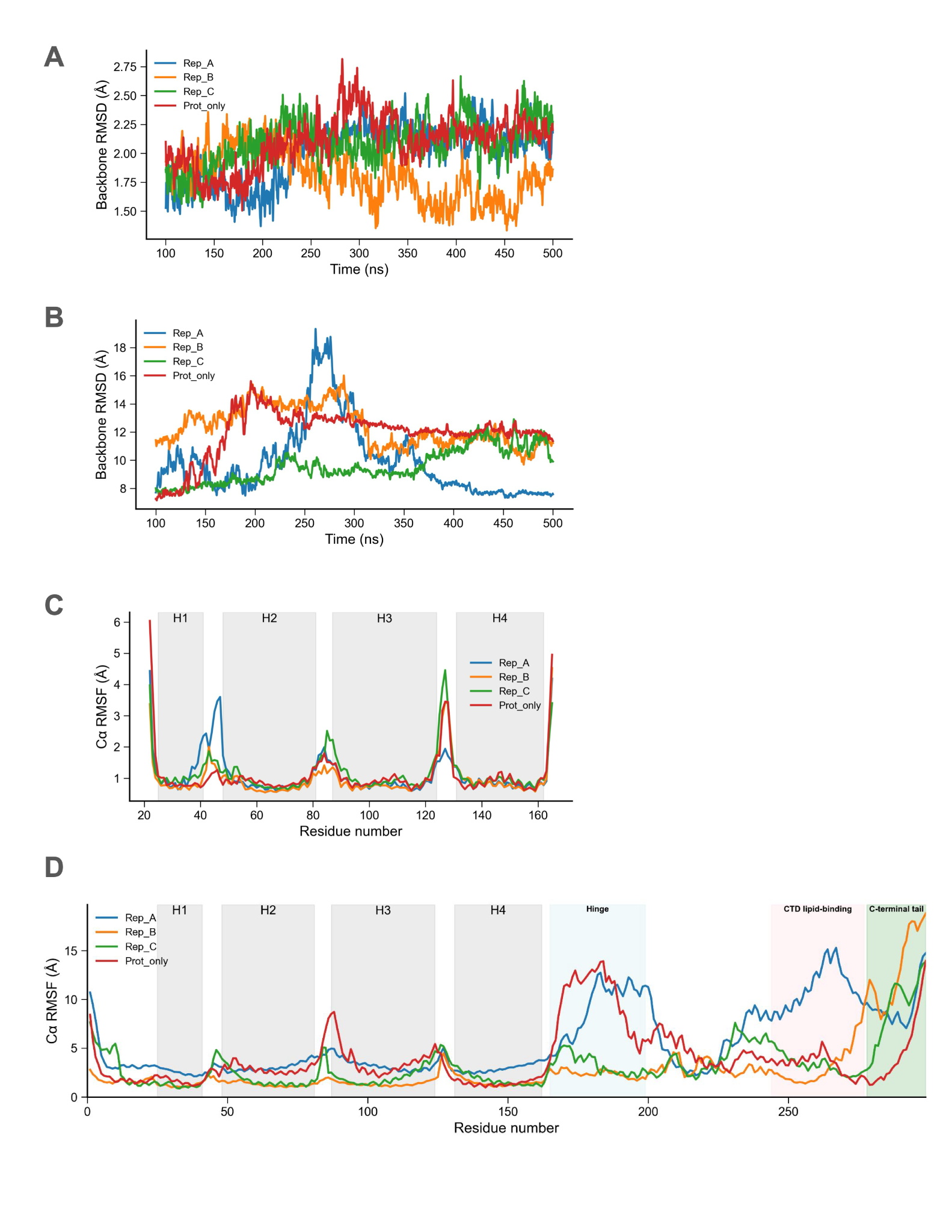

**Fig S2. Equilibration and residue-level flexibility of ApoE4 during molecular dynamics simulations.** **A,B)** Backbone RMSD of the NTD **(A)** and FL ApoE4 **(B)** in membrane-bound simulations (Rep_A, Rep_B, and Rep_C) and in solution-only control simulations lacking a lipid bilayer (Prot_only). **C,D)** Cα RMSF profiles of the NTD **(C)** and FL ApoE4 **(D)** calculated over the production trajectories for the membrane-bound replicates and solution-only controls, highlighting residue-specific differences in conformational flexibility upon membrane association.

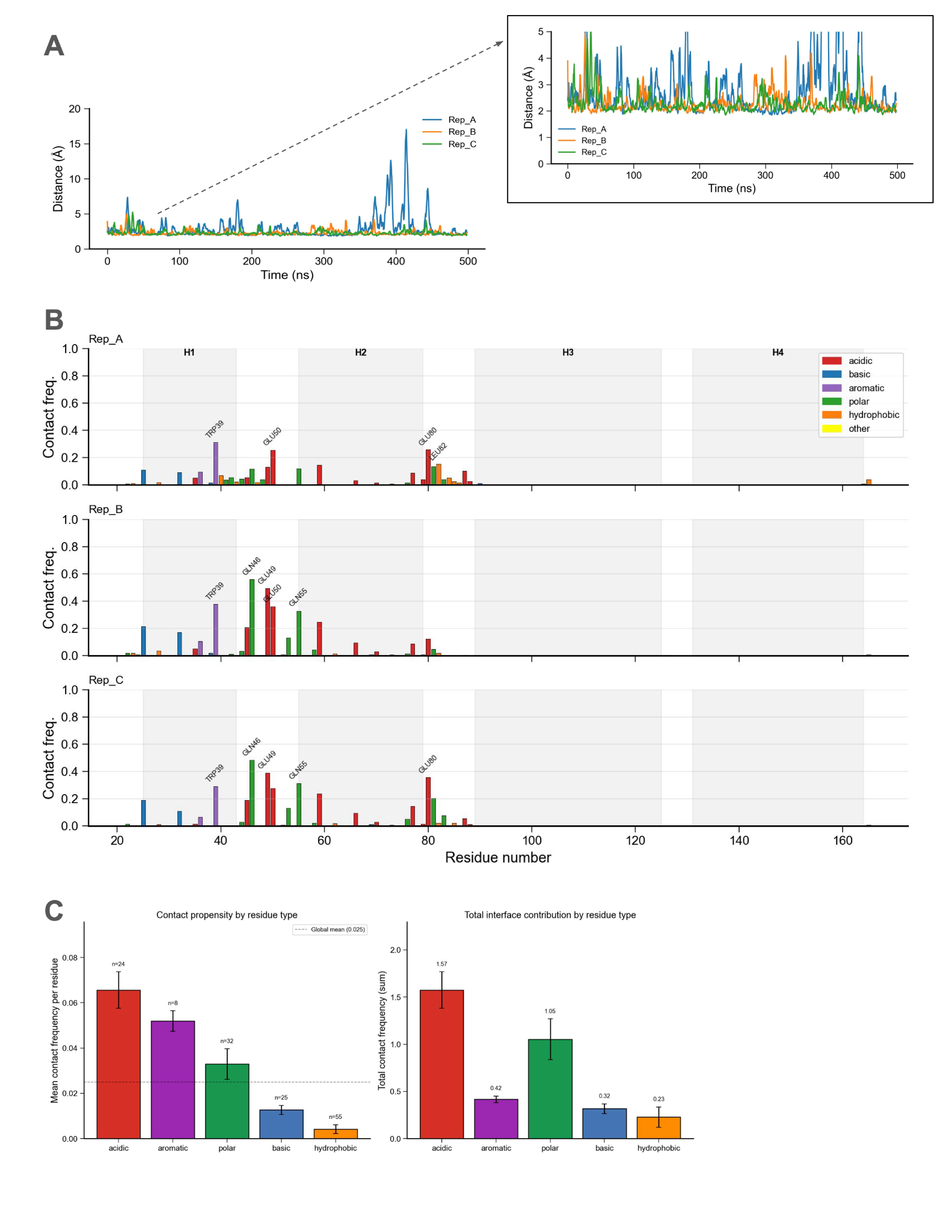

**Fig S3. NTD-DOTAP bilayer contact analysis identifies the H1-H2 face as the primary binding interface. A)** Time-resolved minimum distance between NTD and DOTAP atoms for each of three independent replicates, showing sustained membrane engagement (distances <3.5 Å indicate direct contact). B) Per-residue contact frequency (fraction of frames with any NTD atom within 6 Å of any DOTAP atom) for the three replicates, demonstrating that NTD consistently engages the bilayer through residues on the H1-H2 face. C) Left: mean contact frequency averaged across residues of each type (acidic, basic, polar, hydrophobic). Right: total contact contribution per residue type, computed as the sum of individual residue contact frequencies. Error bars indicate SEM across the three replicates.

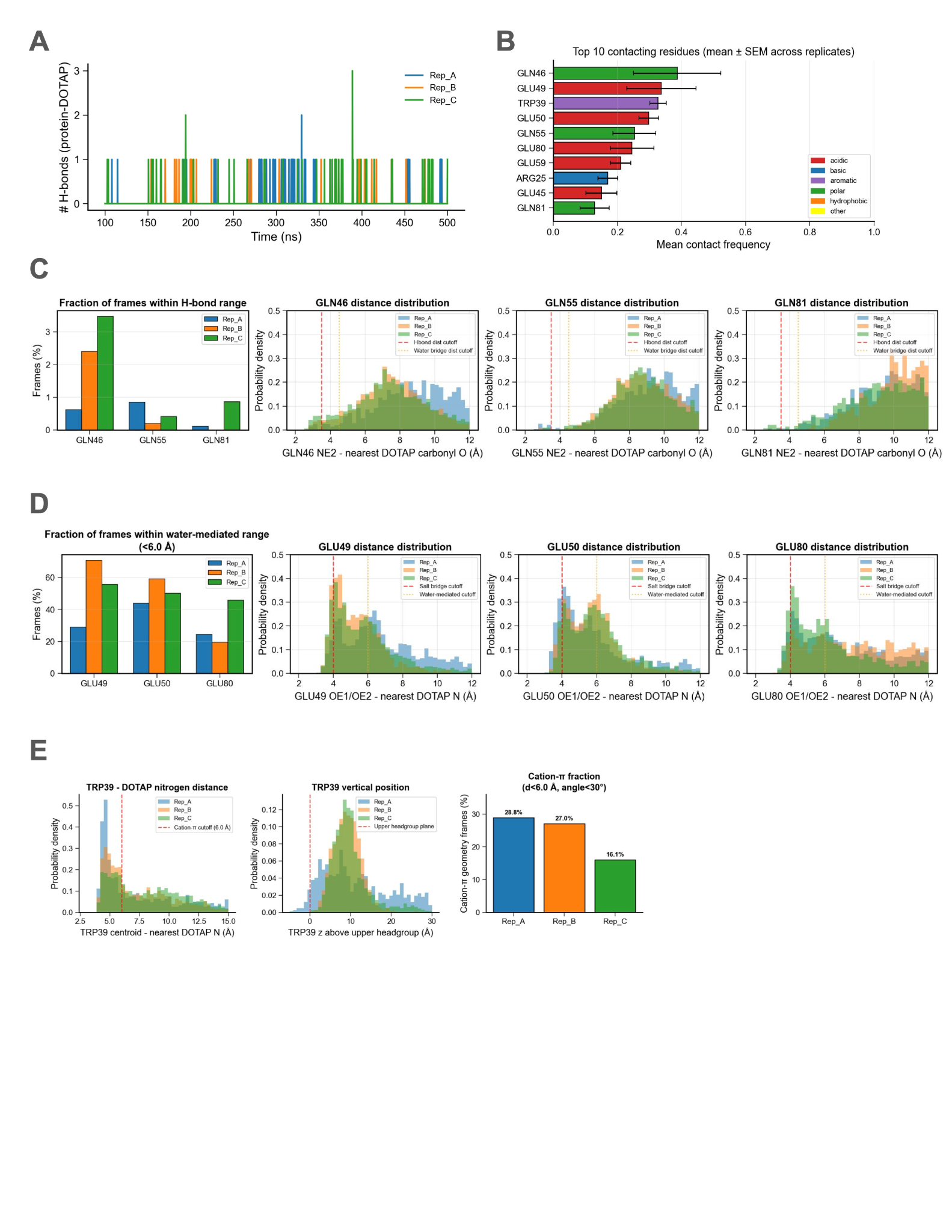

**Fig S4. Detailed analysis of NTD-DOTAP interactions identifies key GLN, GLU, and TRP residues as the dominant contributors to binding. A)** Number of H-bonds between NTD and DOTAP throughout the analysis window (last 400 ns) for each of three replicates, showing limited H-bonding. **B)** Top 10 per-residue contact frequencies averaged across the three replicates, identifying GLN46, GLN55, and GLN81 as the dominant polar residues; GLU49, GLU50, and GLU80 as the dominant acidic residues; and TRP39 as the dominant aromatic residue. Error bars show SEM across the replicates. **C)** Per-residue H-bond analysis for key GLN residues: percentage of frames within H-bond range and corresponding distance distributions (donor-acceptor distance <3.5 Å, angle >120°). **D)** GLU-DOTAP interaction analysis. Left: fraction of frames within water-mediated range (<6.0 Å between GLU OE1/OE2 and the nearest DOTAP nitrogen) for the top three engaging acidic residues. Right: distance distributions for GLU49, GLU50, and GLU80 across three replicates, with vertical lines indicating the salt bridge cutoff (4 Å, red) and water-mediated cutoff (6 Å, orange). The distributions peak in the 4-6 Å range, indicating that GLU residues interact with DOTAP predominantly through water-mediated electrostatics rather than direct salt bridges. **E)** TRP39-DOTAP cation-π interaction analysis. Left: distance distribution between the TRP39 indole ring centroid and the nearest DOTAP trimethylammonium nitrogen across three replicates (peak at 4-5 Å, within the cation-π range). Center: TRP39 ring centroid z-position relative to the upper headgroup plane (TRP39 sits ~10 Å above the headgroup throughout the simulation). Right: fraction of frames in cation-π geometry (centroid distance <6.0 Å, angle between ring normal and N-centroid vector <30°): 28.8%, 27.0%, and 16.1% for Rep_A, Rep_B, and Rep_C respectively. TRP39 anchors the NTD to the bilayer surface through cation-π interactions with DOTAP headgroups.

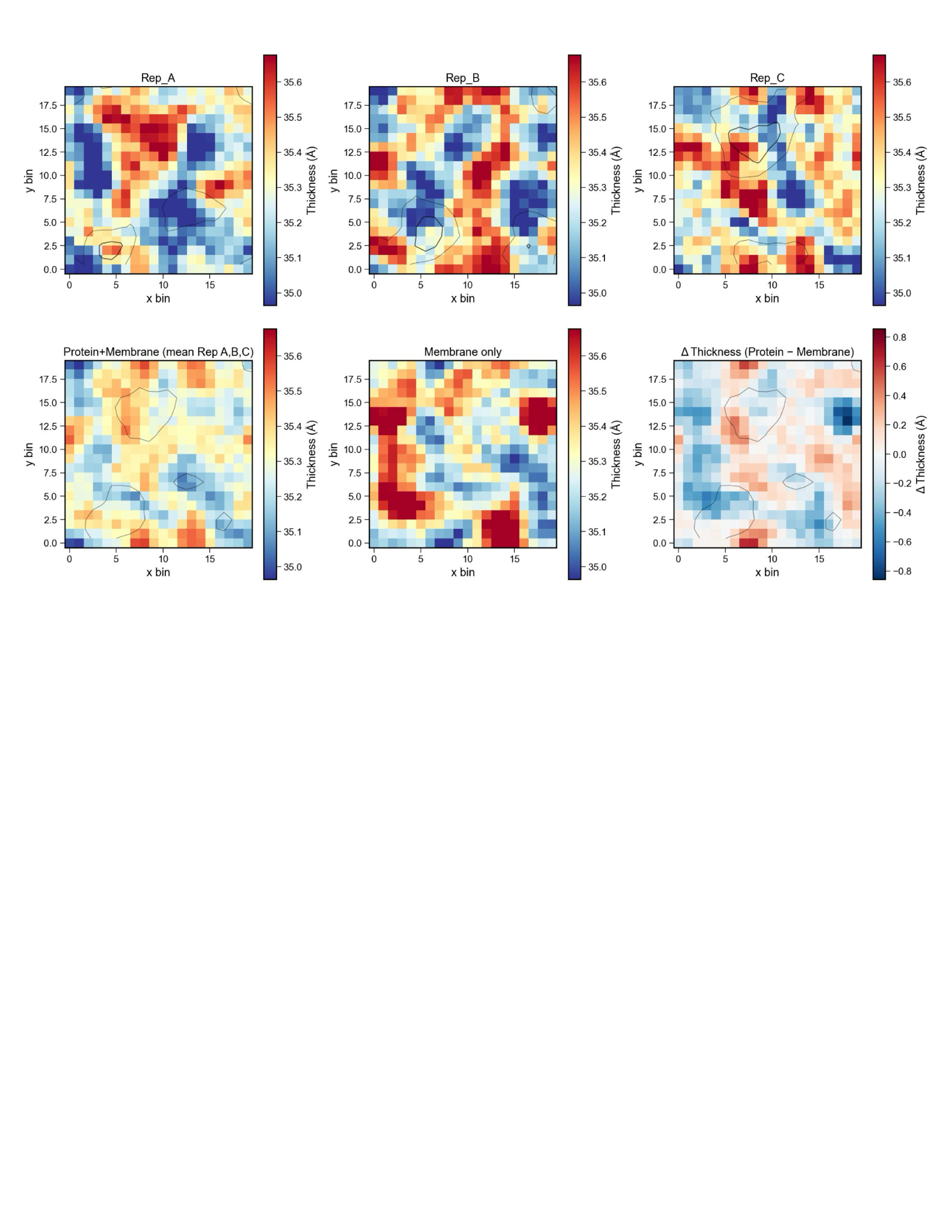

**Fig S5. NTD-induced membrane perturbation results in localized bilayer thinning.** The top row shows the average bilayer thickness map in the presence of the NTD for Rep_A (left), Rep_B (center), and Rep_C (right), calculated over the analysis window (last 400 ns of each trajectory). The projected protein footprint is overlaid to illustrate the spatial relationship between membrane thinning and protein localization. The bottom row presents the mean bilayer thickness map averaged across the three replicates (left), the membrane-only control system (center), and the difference in thickness between the NTD-containing systems and the membrane-only control (right). Regions of reduced thickness are localized near the protein footprint, indicating that NTD binding induces modest membrane thinning without extensive disruption of bilayer structure.

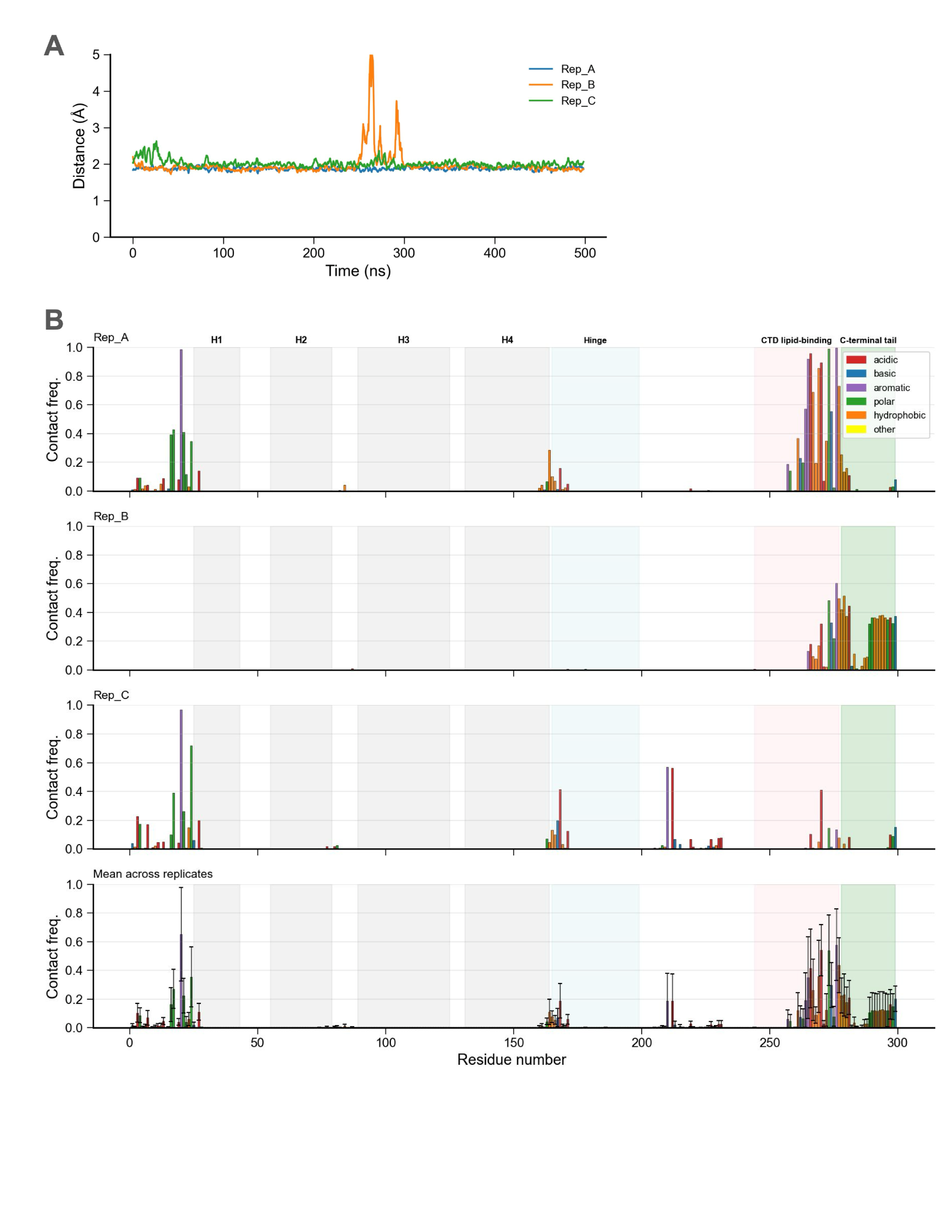

**Fig S6. FL ApoE4–DOTAP bilayer contact analysis reveals three distinct orientation-dependent membrane-binding modes. A)** Time-resolved minimum distance between full-length (FL) ApoE4 and DOTAP atoms for three independent simulation replicates. Distances below 3.5 Å indicate direct protein–lipid contact and demonstrate stable membrane association throughout the simulations. **B)** Per-residue contact frequency, defined as the fraction of simulation frames in which any atom of a given FL ApoE4 residue is within 6 Å of any DOTAP atom. The three replicates exhibit distinct membrane-binding orientations. In Rep_A, membrane contacts are concentrated primarily within the C-terminal domain (CTD) amphipathic helix region, with additional contributions from a subset of N-terminal domain (NTD) residues. In Rep_B, membrane association is dominated by residues at the distal end of the CTD lipid-binding region and the C-terminal tail. In Rep_C, contacts are distributed across residues in the NTD, hinge region, and portions of the CTD, reflecting a more heterogeneous binding mode. The average contact profile across all replicates highlights the CTD lipid-binding region as the principal determinant of FL ApoE4 membrane association.

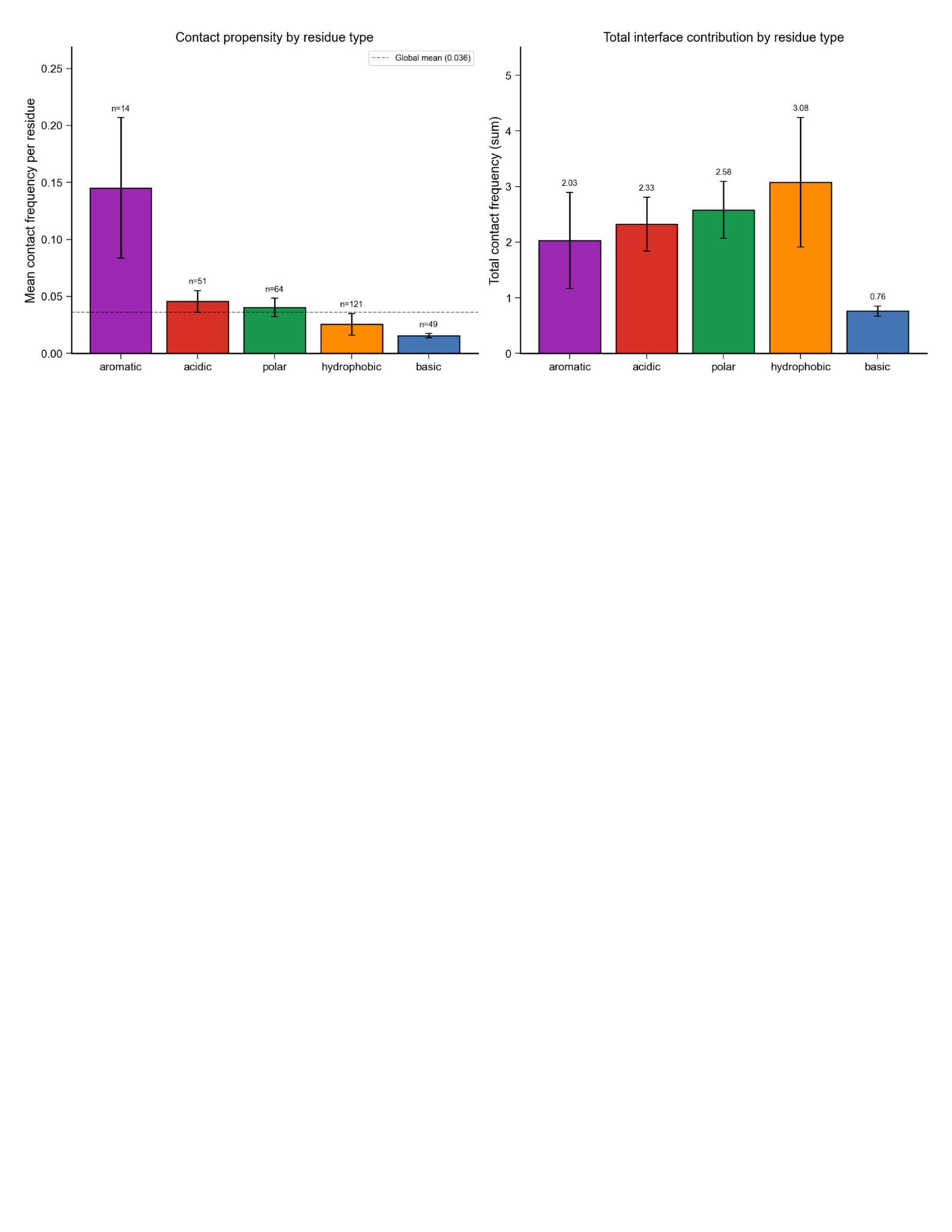

**Fig S7. FL-DOTAP contact propensity by residue type.** Left: mean contact frequency averaged across residues of each type (aromatic, aliphatic hydrophobic, acidic, basic, polar). Right: total contact contribution per residue type, computed as the sum of contact frequencies across all residues of that type. Error bars indicate SEM across the three replicates.

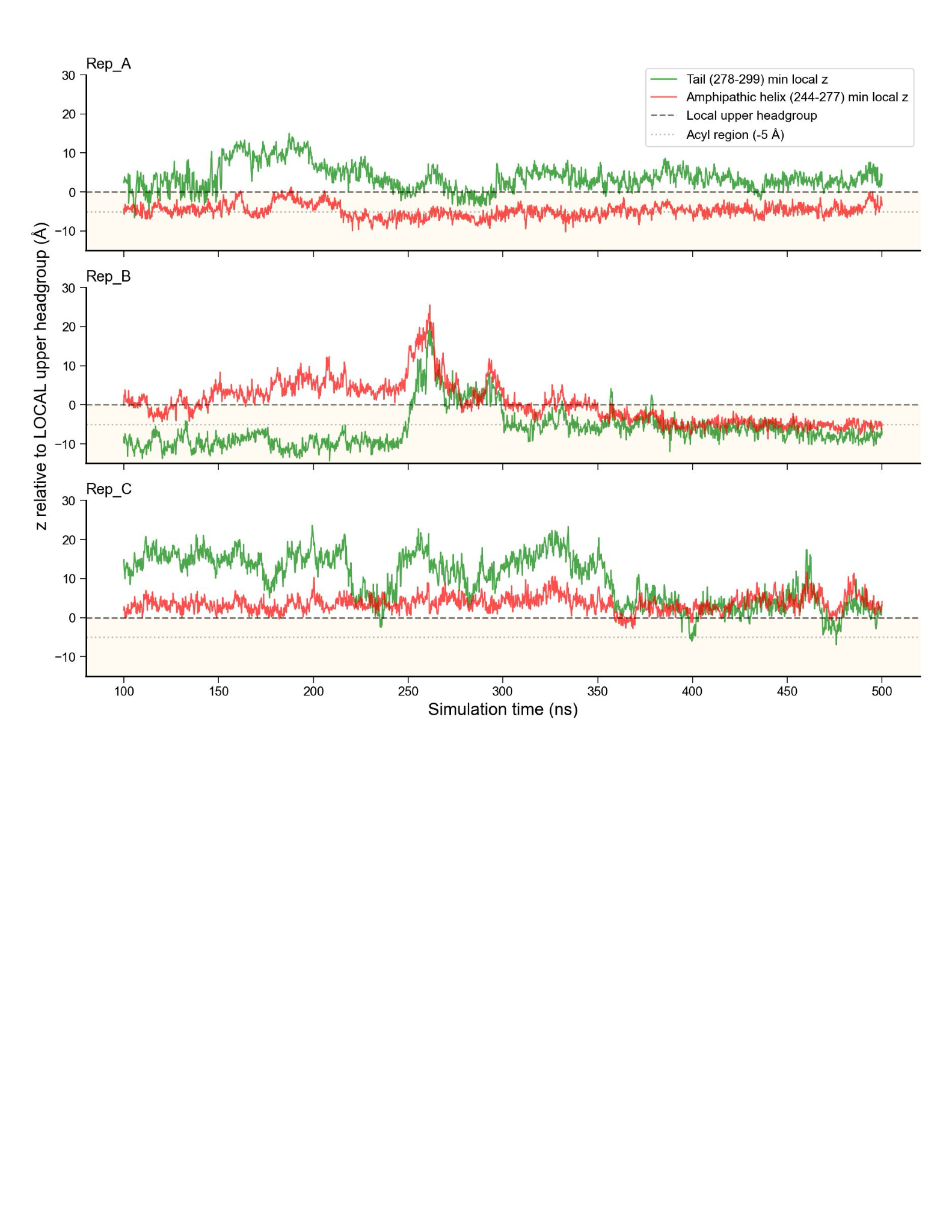

**Figure S8. Time-resolved local-reference depth analysis reveals distinct CTD engagement modes across replicates.** Minimum z-position of the amphipathic helix (red, residues 244-277) and C-terminal tail (green, residues 278-299) relative to the local upper headgroup plane over the analysis window. Top (Rep_A): the amphipathic helix is continuously inserted at the bilayer interface throughout the simulation, while the C-terminal tail remains at or above the headgroup plane. Center (Rep_B): the C-terminal tail inserts deeply into the bilayer with the amphipathic helix remaining in the aqueous phase, until ~250 ns when the tail dissociates from the bilayer. The protein later re-engages with the bilayer; trajectory inspection reveals that while the minimum z-position recovers to comparable values, only a small subset of residues from each region contacts the bilayer in this new configuration, with the bulk of both the helix and tail remaining in the aqueous phase. Bottom (Rep_C): both regions remain above the bilayer throughout the simulation, consistent with the inverted starting orientation preventing sustained CTD engagement.

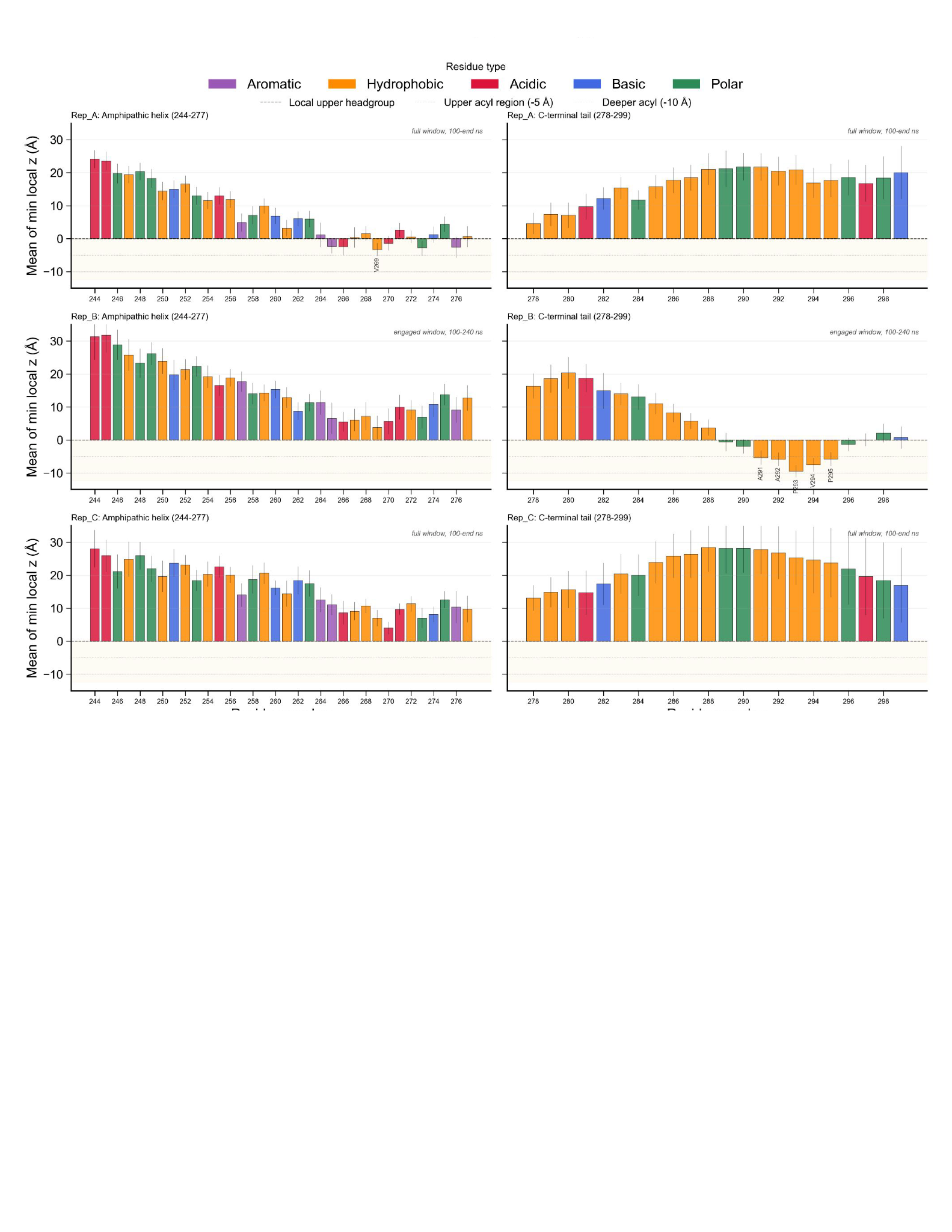

**Figure S9. Per-residue local-reference depth profiles reveal the specific residues anchoring each CTD engagement mode.** Mean of the minimum local z-position per residue in the amphipathic helix (left column, residues 244-277) and C-terminal tail (right column, residues 278-299) for each replicate. Error bars indicate the standard deviation across the analysis window. Residues are color-coded by type: aromatic (purple), aliphatic hydrophobic (orange), acidic (red), basic (blue), polar (green). Reference lines indicate the local upper headgroup plane (dashed, z = 0), upper acyl region (z = -5 Å), and deeper acyl region (z = -10 Å). **Top (Rep_A):** PHE265, GLU266, VAL269, GLN273, and TRP276 are consistently positioned at or below the local headgroup plane in the amphipathic helix, while the C-terminal tail residues remain well above the bilayer. **Middle (Rep_B, engaged window 100-240 ns)**: the inverse pattern is observed, with the amphipathic helix above the bilayer and the C-terminal tail residues 290-296 sustaining deep insertion into the upper acyl region, consistent with the hydrophobic character of residues 291-295 (ALA, ALA, PRO, VAL, PRO). **Bottom (Rep_C):** both regions remain above the bilayer throughout the simulation.

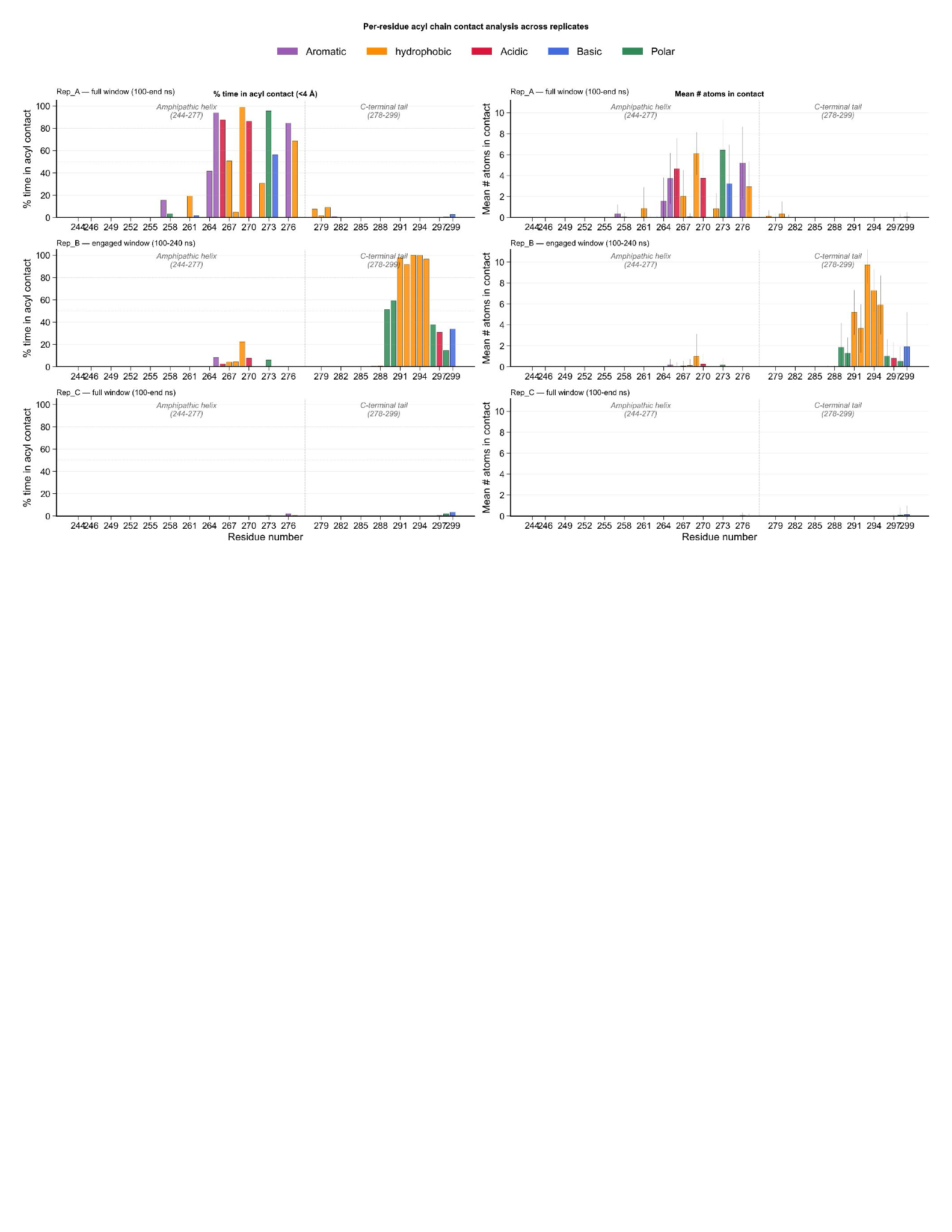

**Figure S10. Per-residue acyl chain contact analysis confirms partial embedding of CTD residues into the bilayer hydrocarbon region.** Two complementary metrics are shown for each residue in the amphipathic helix (residues 244-277) and C-terminal tail (residues 278-299) across all three replicates. **Left column:** percentage of frames in which any residue heavy atom is within 4 Å of any DOTAP acyl chain carbon, computed over the analysis window (last 400 ns for Rep_A and Rep_C; engaged window 100-240 ns for Rep_B). **Right column:** mean number of residue heavy atoms simultaneously in acyl contact per frame; error bars indicate the standard deviation across the analysis window. Residues are color-coded by type: aromatic (purple), aliphatic hydrophobic (orange), acidic (red), basic (blue), polar (green). Together, these metrics identify PHE265, GLU266, VAL269, GLN273, and TRP276 as the dominant amphipathic helix anchors in Rep_A and residues 291-295 (ALA, ALA, PRO, VAL, PRO) as the dominant C-terminal tail anchors in Rep_B. Rep_C shows minimal acyl contact in both regions, consistent with its inverted starting orientation preventing sustained CTD engagement.

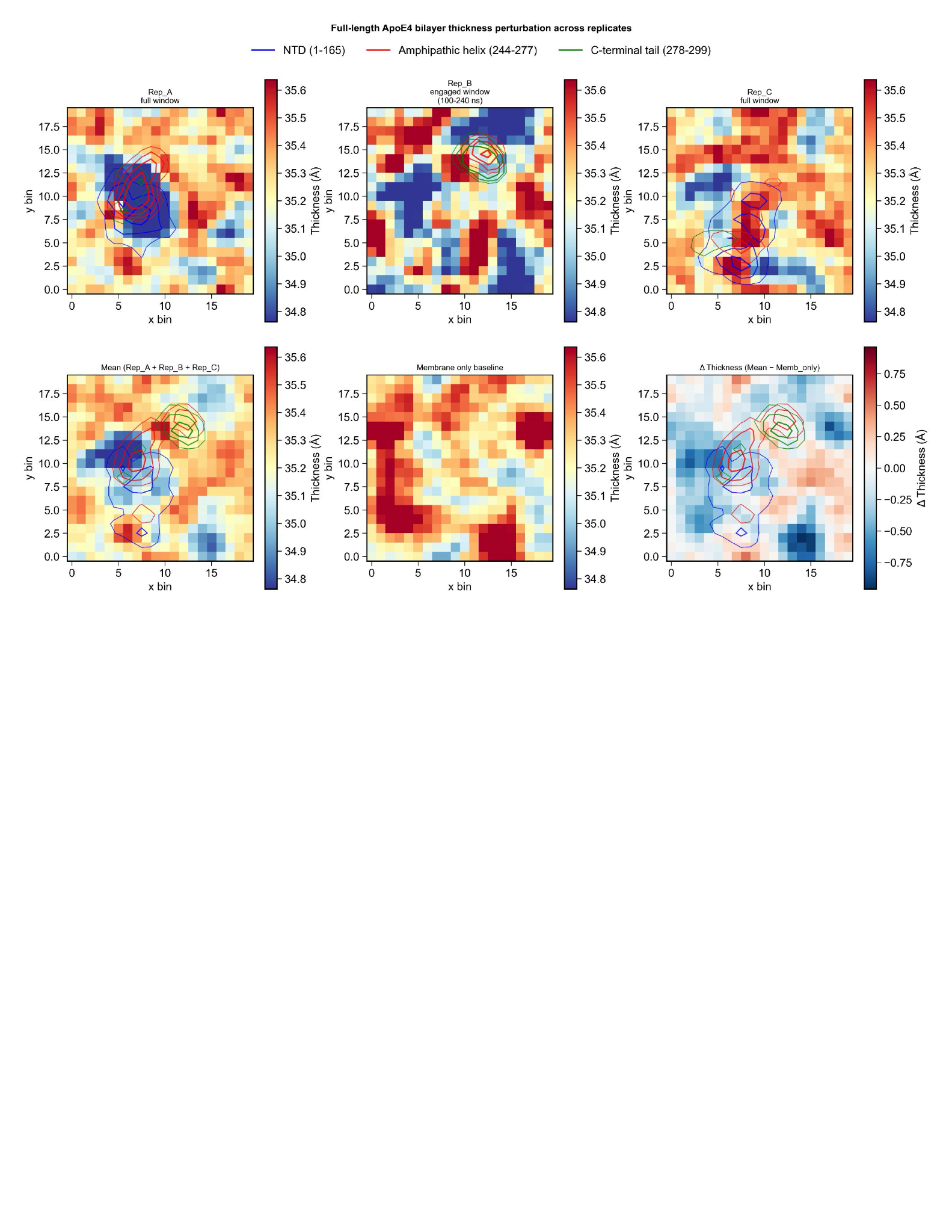

**Figure S11. FL-induced bilayer perturbation shows replicate-specific patterns of localized thinning and thickening.** **Top row:** time-averaged bilayer thickness maps in the presence of full-length ApoE4 for Rep_A (left), Rep_B (center), and Rep_C (right), computed over the analysis window (last 400 ns for Rep_A and Rep_C; engaged window 100-240 ns for Rep_B). Mean protein contact footprints are overlaid as colored contours: NTD (blue), amphipathic helix (red), C-terminal tail (green). **Bottom row:** mean bilayer thickness map averaged across the three replicates (left), the membrane-only control simulation (center), and the Δ thickness map computed as the difference between the mean FL-containing systems and the membrane-only control (right). Rep_A shows localized bilayer thinning beneath the amphipathic helix footprint, consistent with the surface-aligned helix compressing the local upper leaflet. Rep_B shows localized thickening under the C-terminal tail engagement footprint, consistent with the tail diving into the bilayer and anchoring lipid headgroups outward. Rep_C shows modest thickening under the NTD-down orientation.
